# Widespread introgression and a potential role for a neo-sex chromosome in booby diversification and speciation

**DOI:** 10.64898/2026.07.17.739190

**Authors:** Danny Jackson, Erik Funk, Linelle Ann L. Abueg, David J. Anderson, Jennifer Balacco, Giulio Formenti, Nivesh Jain, Vicki L. Friesen, Thomas C. Mathers, James Morris-Pocock, Marco Sollitto, Tammy Steeves, Tatiana Tilley, Vertebrate Genomes Project Consortium Phase I, Melissa Wilson, Carlos Zavalaga, Scott A. Taylor

**Affiliations:** Comparative Genomics and Reproductive Health Section, Center for Genomics and Data Science Research, National Human Genome Research Institute, National Institutes of Health, Bethesda, MD, 20852 USA; University of Colorado, Ecology and Evolutionary Biology; The Vertebrate Genome Lab, The Rockefeller University, New York, NY 10128; Wake Forest University, Biology; Queen’s University, Biology; Tree of Life, Wellcome Sanger Institute, Cambridge, UK; St. Lawrence College, School of Baccalaureate Nursing; Department of Biology, University of Florence, Sesto Fiorentino 50019, Italy; University of Canterbury, School of Biological Sciences; Universidad Cientifica del Sur, Unidad de Investigación en Ecología y Conservación de Aves Marinas

**Keywords:** Hybridization, speciation, seabirds, neo-sex chromosome, evolution, marine

## Abstract

Speciation in marine taxa is dynamic and complex, often occurring in the absence of absolute geographic barriers to gene flow. Boobies comprise a clade of seven highly mobile seabird species with high diversity in the eastern Pacific Ocean, where no land barriers separate extant sister species and hybridization is widespread. They exhibit striking diversity in bare-part coloration, an important signal of mate quality, and possess a novel multiple sex chromosome system (Z_1_Z_2_Z_1_Z_2_ / Z_1_Z_2_W), likely resulting from a fusion between an ancestral autosomal microchromosome and the W chromosome. To understand speciation processes in the context of gene flow, we sequenced and analyzed 29 short-read booby genomes and assembled a reference northern gannet genome to (1) test for introgression, (2) characterize genomic patterns of divergence across species, and (3) investigate the gene content and evolution of the neo-sex chromosome. We found that divergence among eastern Pacific sister taxa is temporally associated with periods of glacial maxima. Between periods of glacial maxima, two pairs of sister species (blue-footed and Peruvian boobies; masked and Nazca boobies) exhibit strong signatures of episodic introgression, indicating that speciation has occurred over multiple periods of divergence followed by gene flow. However, in the third pair of sister species, genomic signatures showed higher divergence within brown booby populations than between brown and Cocos boobies. We also identified the epidermal differentiation complex – a gene cassette involved in skin, feather, beak, and claw development – on the neo-sex region of the W chromosome, where it spans the boundary between the putatively recombining and non-recombining regions, with interspecific variation in the exact recombination suppression boundary. We inferred that this complex may contribute to diversity in feather and bare part coloration across boobies. Together, these results reveal that diversification and speciation in a clade of highly mobile seabirds emerged from a complex evolutionary history involving historical climate dynamics, introgression, and sex chromosome evolution.

## Introduction

Speciation in the absence of geophysical barriers to dispersal depends on mechanisms that maintain divergence despite the presence of gene flow (8, 9). The dynamism and interconnectivity of the world’s oceans foster extraordinary and complex evolutionary processes (10–16). Seabirds are highly mobile but variably philopatric to isolated breeding islands spread throughout vast ocean basins (17), and seabird speciation often occurs in the absence of any absolute geographic barriers to dispersal (18, 19). In these circumstances, speciation mechanisms driven by nongeographical factors or dynamic physical barriers might be expected (12, 20–23). Pantropical species display a distribution pattern that crosses continents, and they breed on isolated islands spread inconsistently throughout enormous ocean basins. Thus the Isthmus of Panama, the African continent, and the vast expanse of Pacific Ocean with few suitable breeding islands serve as major barriers to gene flow (24). Yet growing evidence indicates that seabirds will cross or navigate around these barriers (25), and the mechanisms of speciation in the absence of complete barriers to gene flow are less well understood (25). Uncharacterized historical barriers to dispersal may also have contributed to divergence. For instance, Pleistocene glaciers facilitated speciation in some contemporarily sympatric polar seabird species (18, 26). Non-polar species show evidence of severe population bottlenecks during periods of glacial maxima (27), which suggests that climatic and biogeographic changes associated with global glaciation events may have promoted divergence and speciation events even in tropical species.

Boobies (family Sulidae, genus *Sula*) exhibit several features that make them a compelling system for exploring speciation processes, including widespread hybridization, putative parapatric speciation, and a fusion between the W chromosome and an autosomal microchromosome (28–32), which all have the potential to impact mechanisms of population differentiation. They comprise seven tropical and subtropical species with high species richness in the eastern Pacific (Figure 1) (33, 34). Three species are pantropically distributed: red-footed booby (*Sula sula*), brown booby (*S. leucogaster*), and masked boobies (*S. dactylatra*). The other four species are endemic to the eastern Pacific Ocean: Nazca booby (*S. granti*), blue-footed booby (*S. nebouxii*), Peruvian booby (*S. variegata*), and Cocos booby (*S. brewsteri*). One pair of sister species, blue-footed and Peruvian boobies, are both endemic to the Eastern Pacific and are allopatric except for a hybrid zone off the coast of northern Peru. The other two eastern Pacific species were considered subpopulations of pantropical species but both were split within the last 30 years: Nazca from masked boobies (32, 35) and Cocos from brown boobies (36, 37). Thus, species diversity in this clade is uniquely rich in the eastern tropical and subtropical Pacific. The processes that shaped speciation and diversification of boobies may illuminate mechanisms of evolution that are important for highly mobile organisms that depend on the world’s oceans.

**Figure 1:**
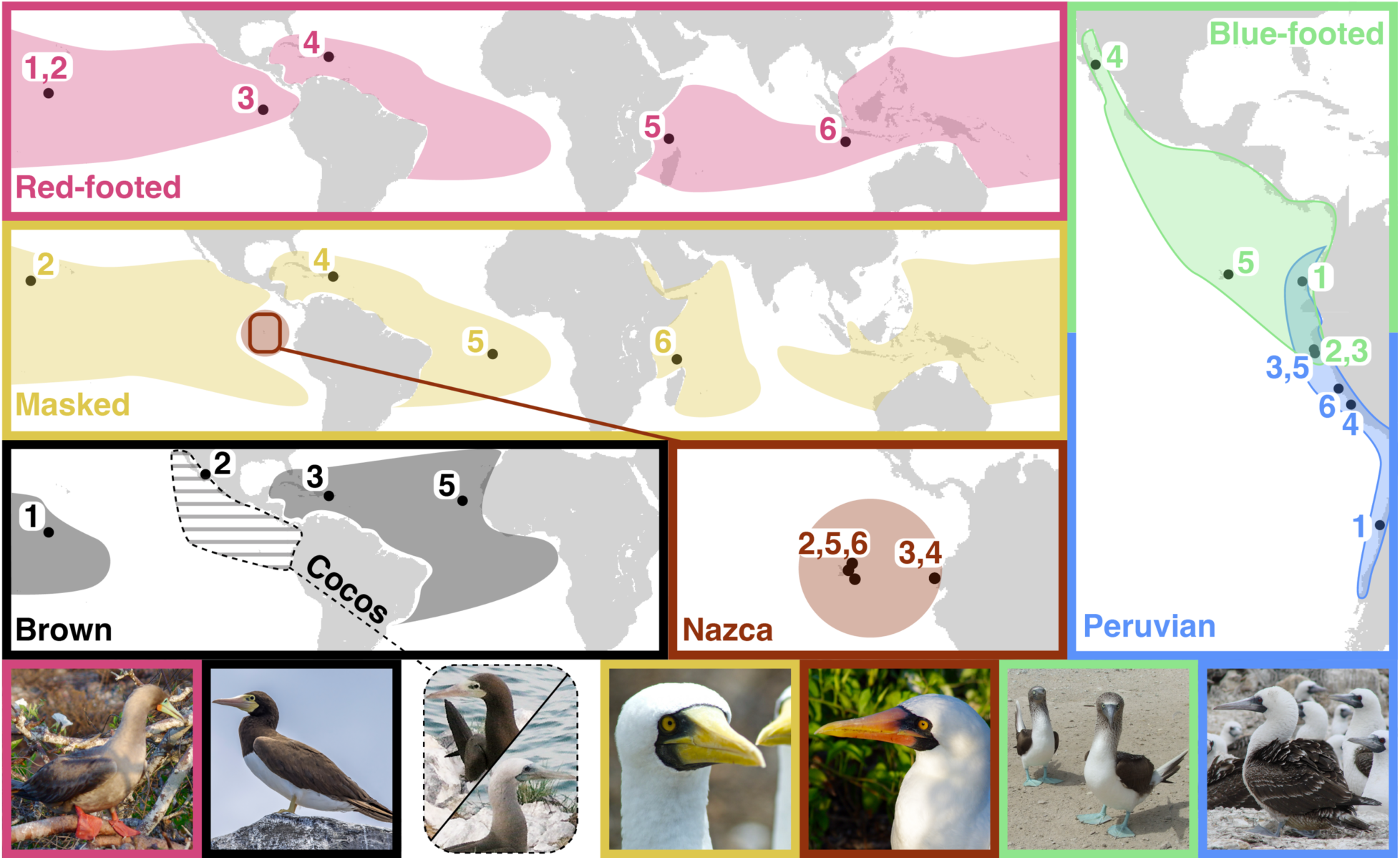
Species breeding ranges, sampling locations and species identity of each individual genome. Sample locations are as follows: **blue-footed booby**: 1. La Plata, 2 & 3. Lobos de Tierra, 4. Isla San Ildefonso, 5. Seymour Norte; **Peruvian booby**: 1. Isla Pajaros, 3. Lobos de Tierra, 4. North Chincha, 5. Lobos de Afuera, 6. Mazorca Island; **red-footed booby**: 1 & 2. Palmyra Atoll, 3. Genovesa, 4. Monito Island, 5. Aldabra Atoll, 6. Christmas Island; **brown booby**: 1. Palmyra Atoll, 2. (***Cocos booby***) Farallon de San Ignacio, 3. Monito Island, 5. Cape Verde; **masked booby**: 2. Johnston Atoll, 4. Monito Island, 5. Ascension Island, 6. Cosmoledo Atoll**; Nazca Booby**: 2. Espanola, 3 & 4. La Plata, 5. Daphne, 6. Genovesa. Species breeding ranges are based on maps from Birds of the World (110–116) and Nelson 1978 (34). Two photographs were included to demonstrate phenotypic similarities and distinctions between brown and Cocos boobies. While female Cocos boobies (top) appear similar to brown boobies, male Cocos boobies (bottom) have a white head and blue face that is distinct from brown boobies. Most photos were taken by authors: Scott Taylor (red-footed, Nazca, blue-footed, and Peruvian boobies) and Tammy Steeves (masked booby). The remaining photos were sourced from Wikimedia commons: Charles J. Sharp (brown booby, https://commons.wikimedia.org/w/index.php?curid=112750434) and Cozcacua (Cocos boobies, https://commons.wikimedia.org/w/index.php?curid=33045834).

The persistence of species differences in the presence of hybridization indicates that genotypic incompatibilities have accumulated between species but are not yet sufficient to prevent interbreeding (8). Morphological evidence suggests that hybridization between booby species is widespread but genetic evidence of hybridization exists for only two species pairs: blue-footed and Peruvian boobies, which hybridize in a narrow region of overlap off the coast of Peru(38), and brown and blue-footed boobies in the Gulf of California (39). Several additional instances of hybridization have been recorded but lack genetic confirmation: Nazca and Cocos boobies both appear to hybridize with their sister taxa, masked and brown boobies, respectively (35, 37); putative hybrids have been observed between masked and brown boobies (33); and interspecies courting and pairing has been documented between Nazca and blue-footed boobies (40). Hybridization alone does not indicate the presence of gene flow between species (introgression), as hybrids can be sterile, not survive to reproductive age, or not successfully breed (8, 9). Introgression has only been documented between blue-footed and Peruvian boobies (31), with no tests applied to other taxa. It is unknown if hybridization acts as a source of genetic variation between diverged species via introgression and, if so, whether it does so in all or in only some instances of documented hybridization.

Non-recombining structural genomic variants can facilitate speciation in the presence of gene flow (41, 42). Recent karyotypic studies of three booby species (red-footed, brown, and masked) found evidence of a neo-sex chromosome, which likely derived from the fusion of a small autosomal chromosome with the W chromosome (28). This led to a Z_1_Z_1_Z_2_Z_2_ / Z_1_Z_2_W sex chromosomal system in boobies, with the former autosomal chromosome inherited in a Z_2_ pattern. The northern gannet (*Morus bassanus*), from the sister genus to boobies, has a typical ZW chromosomal system based on karyotypic analysis (43), therefore the neo-sex chromosome appears synapomorphic to boobies. According to Haldane’s Rule, genes on the sex chromosomes are less likely to introgress and can play a large role in maintaining species boundaries (44). All observed hybrids between Peruvian and blue-footed boobies are parented by a female Peruvian and male blue-footed booby (38), suggesting that variants on the non-recombining regions of the blue-footed booby W chromosome cannot introgress into Peruvian boobies. Additionally, hybrid females have only been observed in mate pairs with male blue-footed boobies (45). The genes within the neo-sex chromosome and interspecific diversity in the structural arrangement of this genomic element are unknown but could facilitate speciation in the absence of physical barriers to gene flow.

Here we address four major evolutionary questions about boobies related to speciation. 1. Do glacial periods correspond to periods of divergence in two sister species pairs of booby? 2. Do species pairs in the Eastern Pacific that demonstrate hybridization also demonstrate introgression? 3. What morphological and physiological processes are associated with genetic variants underlying species differences between recently diverged taxa? 4. What is the gene content of the neo-sex region of the W chromosome, and to what extent has recombination been suppressed in the neo-sex region? We generated and analyzed short-read genome sequences for representatives of all currently recognized species of boobies. Using these genome sequences, we reconstructed the phylogeny of boobies and found a lack of phylogenetic evidence for one species pair that shows evidence of assortative mating, Cocos and brown boobies. We then modeled the demographic histories of booby species, finding strong evidence that glacial periods are associated with periods of divergence in the eastern Pacific Ocean, with interglacial periods of gene flow. We used our phylogeny to test for evidence of introgression, finding evidence of introgression between blue-footed and Peruvian boobies and between masked and Nazca boobies. We identified genes in differentiated genomic regions, finding that genes associated with morphological development are potential drivers of booby diversification. Finally, we identified the ortholog to the neo-Z_2_ chromosome in the northern gannet genome, characterized which species the neo-sex chromosome fusion occurs in, estimated and compared chromosome-wide signatures of recombination suppression across species, and identified the gene content of each region. We found that the epidermal differentiation complex spans the recombination boundary on the W chromosome, with interspecific differences, and we identify this neo-sex chromosome as a putatively important genomic region in maintaining species boundaries in boobies.

### Northern Gannet Genome Sequencing and Assembly

We assembled the first chromosome-level genome for a sulid species from a male northern gannet at the Leibniz Institute for Zoo and Wildlife Research, collected by Jorns Fickel and coordinated by Tom Gilbert. Gannets are the sister genus to boobies, ∼17.2 million years diverged (46). In the context of putative introgression and without any prior knowledge on the intra-genus diversity of the neo-sex chromosome, the use of a gannet reference genome facilitates an analysis of diversity across this chromosome in boobies without in-group bias (47, 48). The genome assembled to 1.3 Gb at 38X coverage with 668 contigs (Contig N50 = 10.2Mb), which were assembled into 34 chromosomes, an assembled mitochondrial genome, and 420 scaffolds.

### Booby Short-Read Genomic Data and Variant Calling

We generated resequencing data from 34 booby individuals, spanning the breeding distributions of all seven species, using samples previously collected for population genetic analyses that were archived at Queens University (Figure 1; Friesen et al. 2002, Morris-Pocock et al. 2011, 2016, Steeves et al. 2003, 2005a, 2005b, Taylor et al. 2010a, 2011a, 2011b). Raw sequence data are publicly available for download through the Sequence Read Archive (BioProject accession: PRJNA836623). We aligned trimmed fastq files to the northern gannet genome that we assembled using bwa mem v0.7.18, excluding four individuals with <3X mean depth of coverage and one with extremely high standard deviation in coverage (Table S1). After filtering, we retained 9,689,372 sites for analyses that could be performed in ANGSD using genotype likelihoods (Principal Component Analyses, and F_ST_) and 17,036,927 sites for all other analyses. Specific filtering parameters are described in the Methods.

### Phylogenetic Relationships and Population Structure

To infer phylogenetic relationships among species and population structure within species, we analyzed autosomal chromosomes using three methods: 1. raxml-ng, 2. principal component analyses (PCA), and 3. ADMIXTURE. We performed PCA and ADMIXTURE analyses (implemented in PCAngsd v1.36.4 (49)) on all individuals, between each pair of sister species, and within each species.

Both approaches produced the same species level relationships identified in previous work (Figures 2, S1, and S2; (50, 51)). Importantly, our findings reinforce previous genetic work and do not support splitting the Cocos booby from the brown booby (2). Brown boobies were paraphyletic with Cocos boobies in all analyses, and we refer to this clade throughout the remainder of the manuscript as the brown-Cocos clade. Red-footed boobies were recovered as sister to all other taxa, with brown and Cocos boobies forming a clade sister to the remaining four taxa. A four-taxon clade is sister to the brown-Cocos clade, with two pairs of sister species within it: blue-footed with Peruvian boobies and masked with Nazca boobies. The PCA and RAxML analyses differed in only a few notable instances (Figures 2, S1, S2, S3). In clustering analyses that included all samples, brown, Cocos, and red-footed boobies cluster together in the plot of PC1 and PC2, whereas the RAxML tree suggests that the brown-Cocos clade should be more similar to blue-footed, Peruvian, masked, and Nazca boobies. Additionally, the admixture analysis of all taxa did not find evidence for either a split between blue-footed and Peruvian boobies or one between masked and Nazca Boobies (Figure S2). However, these splits were recovered in analyses that only included samples within a species pair (Figure S3). F_ST_ comparisons of sister taxa showed slightly more divergence between blue-footed and Peruvian boobies (weighted F_ST_ = 0.222), than between masked and Nazca boobies (weighted F_ST_ = 0.189).

**Figure 2.**
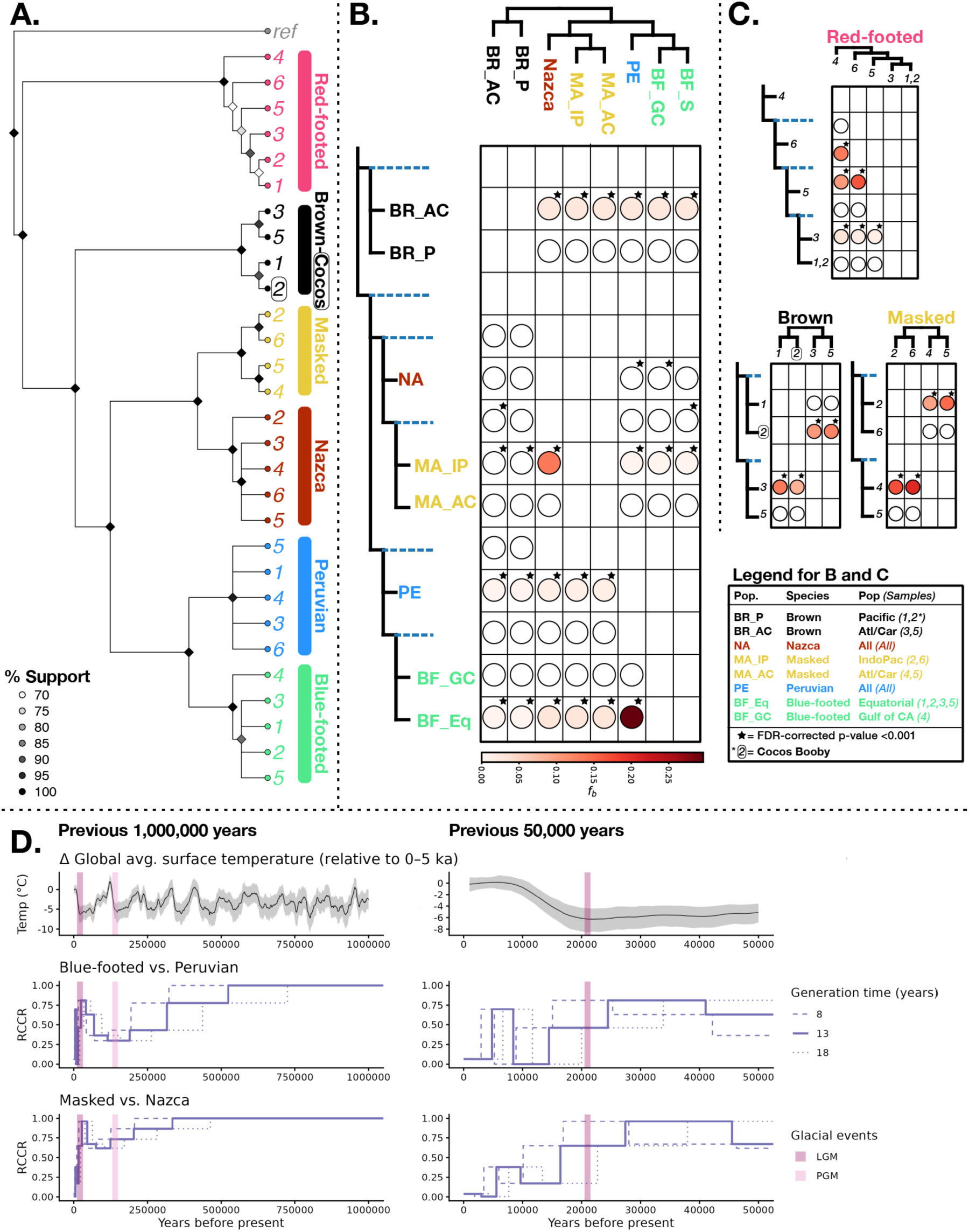
Phylogenetic inference and evidence of introgression. **A.** The consensus tree of the 32 chromosome-wide RAxML best trees. Nodes are colored by the percentage of trees that support the split and nodes with less than 70% support were collapsed. The northern gannet reference genome was used to root the tree (“ref”). **B.** Branch-specific *f_b_* statistics, which display an excess of shared derived alleles between the branch on the y-axis and the population on the x-axis. Significant values are noted with a star. A number of pairs had significantly positive values with very low (<0.001) *f_b_* statistics. **C.** Intraspecific branch-specific *f_b_* statistics, evaluating within-species patterns of gene flow across ocean basins for the three pantropical species: red-footed, brown, and masked boobies. **D.** Relative cross-coalescent plots comparing blue-footed vs. Peruvian boobies and masked vs. Nazca boobies, alongside global change in surface temperatures modeled by Snyder 2016 (105). Pink lines highlight the approximate timing of the last glacial maximum (LGM, ∼21kya) and the penultimate glacial maximum (PGM, ∼140kya) (117). The range of estimated generation times of all boobies are plotted. Generation time estimates are 16.3 and 9.8 years for masked and Nazca boobies, respectively, and are 8.5 and 10.6 years for blue-footed and Peruvian boobies, respectively (118).

We recovered evidence of population structure within all three pantropical species: masked boobies, red-footed boobies, and the brown-Cocos clade (Figures S2-S3). In each case, the structure corresponds to two major land barriers separating tropical oceans: the American continents and the continent of Africa (Figures S2 and S3). Masked boobies split into an Atlantic Ocean/Caribbean Sea clade and a Pacific/Indian Ocean clade. The brown-Cocos clade followed a similar pattern, but we lacked samples from the Indian Ocean, a region we did not sample for brown boobies. Red-footed boobies demonstrate more within-species structure in the Atlantic and Indian Oceans, as the Caribbean Sea sample exhibiting the most diverged signal, followed by the eastern, then western, Indian Ocean (with low support), leaving a monophyletic Pacific Ocean clade. In eastern Pacific species, only blue-footed boobies demonstrated any evidence of population structure, with a split between an individual in the Gulf of California and the samples from the equatorial population. Peruvian and Nazca Boobies showed no within-species geographic clustering patterns (Figures 2A, S2, and S3). These results are consistent with a role for major physical barriers to dispersal in speciation.

### Demographic History Reconstructions

We assessed and found evidence of speciation across potential periodic divergence events followed by secondary contact with gene flow. We modeled relative cross-coalescence rates (RCCR) for two pairs of sister species (blue-footed and Peruvian boobies; masked and Nazca boobies) using MSMC2 (Figure S4), and plotted RCCRs alongside average change in global surface temperature. Specifically, the RCCR plots suggested that the initial stages of speciation occurred during glacial maxima (Figure 2D). Both species pairs showed evidence of a single ancestral population that went through two subsequent periods of divergence followed by secondary contact and gene flow. The two divergence periods temporally align with global temperature patterns, with the penultimate glacial maximum (PGM, ∼140kya) aligning with the deeper divergence period and with the last glacial maximum (LGM, ∼21kya) immediately preceding the more recent divergence period. Both divergence periods demonstrated lower RCCR values indicative of deeper fission between blue-footed and Peruvian boobies than between masked and Nazca boobies. However, the recent signature of gene flow following secondary contact, inferred through higher peak RCCR values, is higher in blue-footed and Peruvian boobies than in masked and Nazca (Figure 2D). We further characterized this signal with direct tests of introgression.

Our RCCR analyses suggested that contemporarily sympatric booby species were likely allopatric during the penultimate and last glacial maxima. Thus, recent speciation in the eastern Pacific arguably was driven by pulsing periods of allopatry followed by secondary contact and gene flow (52). The distribution of Peruvian boobies likely tracked the Humboldt current, but we lack clear predictions of historical distributions of blue-footed, masked, and Nazca boobies. Further investigations into the historic drivers of allopatry in contemporarily sympatric boobies would advance our understanding of seabird speciation processes.

### Introgression Across Species Boundaries

We tested for signatures of introgression across the booby phylogeny based on widespread evidence of hybridization between multiple species. We computed f-branch (*f_b_*) and Patterson’s *D*-statistics using Phred-scaled genotype likelihoods in Dsuite. We found evidence for introgression between multiple booby species pairs and across land barriers within all pantropical booby species. Two pairs of sister species – blue-footed and Peruvian; masked and Nazca – exhibit significant genomic evidence of introgression between sympatric populations in the eastern Pacific Ocean (*f*_b_ > 0.05; Figure 2B). Despite introgression, we observed strong signals of phylogenetic divergence between each pair of sister taxa (Figures 2A, S2); thus, our data show that gene flow has not eroded species differences between either blue-footed and Peruvian or masked and Nazca boobies. These results contrast with the third sister pair, Cocos and brown boobies, which does not show evidence of differentiation consistent with species-level separation (Figure 2A, S2, S3). Instead, the greatest signature of population structure in this clade is between the Pacific and Atlantic/Caribbean populations rather than eastern and Central Pacific Ocean populations (Figure S2, S3). Of note, the Cocos booby showed evidence of gene flow with the brown booby sample from the Caribbean Sea (Figure 2C). Introgression is widespread in birds, and can be observed intermittently throughout a phylogeny (53). However, it is remarkable that boobies show evidence of introgression between all sister taxa. Natural selection may play a large role in reinforcing species boundaries and limiting introgression. Blue-footed and Peruvian boobies appear to be under selection for divergent phenotypes and behaviors in regions of overlap, as multiple foraging-related traits show evidence of greater character displacement in sympatry than in allopatry (54). We further investigate the genomic regions involved in maintaining species boundaries below.

Only one clustering-robust D-statistic was significant after correction for false discovery rate, indicating introgression between masked and Nazca boobies (D= 0.14, Z = 24.94, FDR corrected clustering-robust p-value = 0.013). In windowed analyses of D_fM_ statistics, a wide window on chromosome 2 shows a signature of introgression and may drive this result (annotated as blue points in Figure 3). We found no significantly overrepresented GO terms in regions of the genome that demonstrate signatures of introgression between blue-footed and Peruvian boobies nor between masked and Nazca boobies.

**Figure 3.**
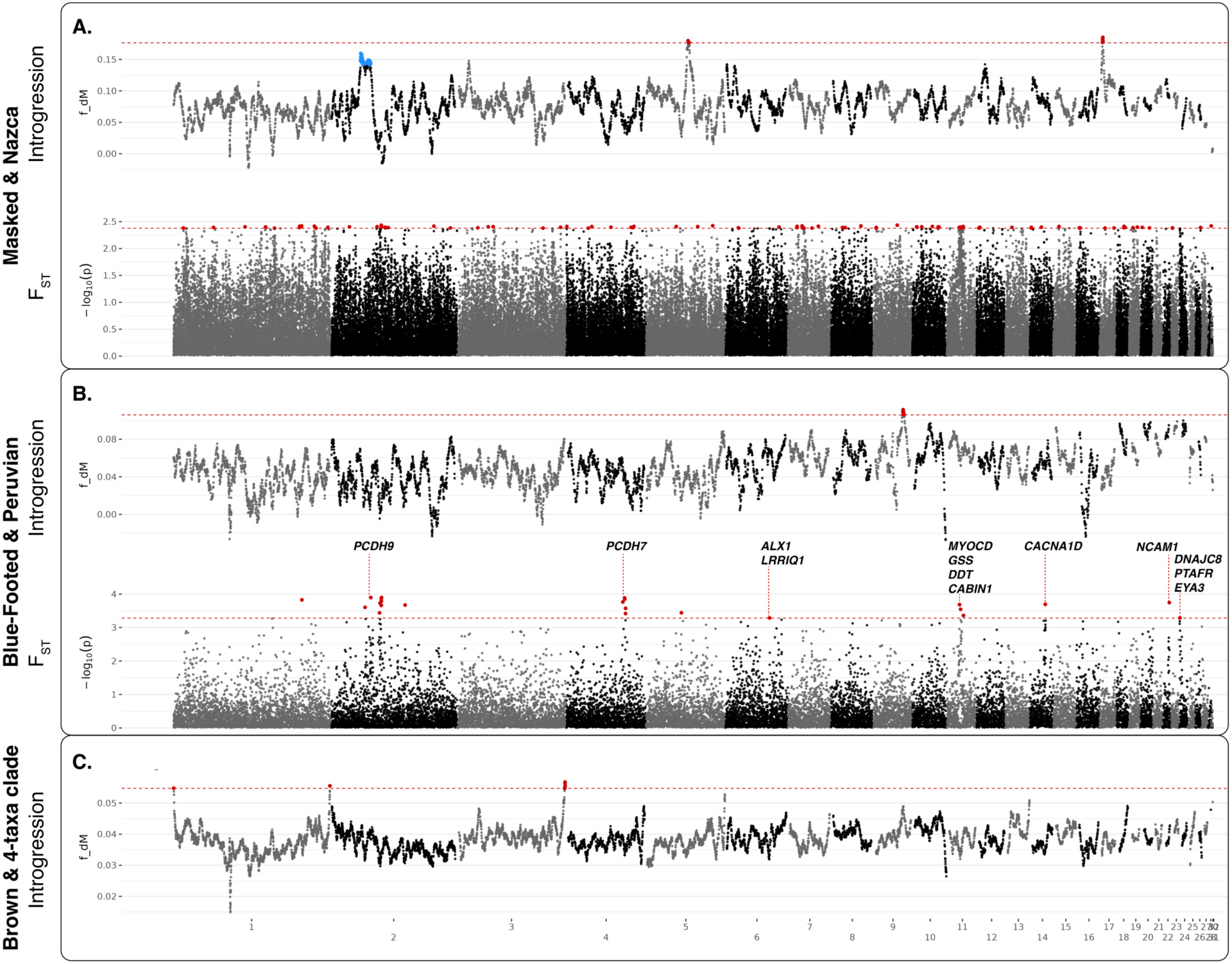
Genome-wide signatures of introgression (*f*_dM_) and differentiation (F_ST_) for three comparisons of interest. In all plots, putative outliers are highlighted as red points, and a dashed red line is used to demarcate the outlier cutoff value. Plot **A** (above) evaluates introgression patterns among sympatric masked and Nazca boobies using population assignments of 1. Atlantic-Caribbean and 2. Indopacific samples of masked boobies and 3. all Nazca boobies. Here, we also show a wide but flat signature of introgression on Chromosome 2, with the top 0.5% windows highlighted in blue. It did not reach our threshold for outliers, but we note it due to its wide signature. Plot **A** (below) evaluates differentiation using all samples of masked boobies and all samples of Nazca boobies. Plot **B** (above) evaluates introgression patterns among sympatric blue-footed and Peruvian booby populations, using population assignments of 1. Gulf of California and 2. equatorial blue-footed boobies and 3. all Peruvian booby samples. Plot **B** (below) evaluates differentiation using all samples of blue-footed boobies and all samples of Peruvian boobies. Plot **C** evaluates introgression between Atlantic-Caribbean brown boobies and the 4-taxon clade (masked, Nazca, blue-footed, and Peruvian boobies), using population assignments of 1. Pacific Ocean and 2. Atlantic-Caribbean brown boobies, and 3. all samples of the 4-taxon clade. Red-footed boobies were used as the outgroup in all introgression comparisons. All plots use a window size of 50kb, and genes within 10kb on either side of the top 0.1% of windows are annotated on discernible peaks. Elevated F_ST_ signals were diffuse for masked and Nazca comparisons, so we did not annotate genes on any peaks. The full list of genes in the outlying regions can be found in the Supporting Dataset.

We also found an unexpected pattern of introgression between the Atlantic/Caribbean population of brown boobies and the four-taxon clade that contains masked, Nazca, Peruvian, and blue-footed boobies. This test was significant regardless of whether these four species were considered individually or treated as a single clade. This contrasts with our hypotheses: based on evidence of hybridization between Cocos and blue-footed boobies in the eastern Pacific, we expected to observe introgression between Pacific brown boobies and blue-footed boobies. This signal of introgression could be caused by ancestral population structure or introgression with an extinct booby species in the Atlantic Ocean or Caribbean Sea — indeed, a diversity of *Sula* fossils from extinct species have been found throughout the tropics (55). The cause of this signature of introgression is conjecture, yet it contributes the important context that, while contemporary species diversity is concentrated to the eastern Pacific, temporally dynamic evolutionary forces have shaped booby diversity throughout the globe. Future analyses with deeper sampling within colonies of brown boobies may help to resolve this unusual signal.

Additional evidence indicated significant but low levels of introgression between many sets of taxa, which we consider to be spurious results driven by correlated *f_b_* statistics resulting from instances of introgression. For example, equatorial blue-footed boobies demonstrate significant signatures of introgression with all other tested species, but the *f_b_* statistic is highest in the comparison with Peruvian boobies and decreases with phylogenetic distance. Intraspecific analyses — including of the brown-Cocos clade — showed evidence of gene flow between all populations of pantropical species.

The eastern Pacific represents a unique region of seabird endemism and diversification. This region is home to multiple sister species pairs and genetically differentiated populations across seabird lineages, including three sister pairs of storm-petrel species (*Hydrobatinae*) (56); distinct populations of brown pelicans (*Pelecanus occidentalis*) (57), magnificent frigatebirds (*Fregata magnificens*) (58, 59), and great frigatebirds (*F. minor*) (60, 61); and several species endemic to the Galápagos Islands and surrounding area, including flightless cormorants (*Phalacrocorax harrisi*) (62), Nazca boobies (*Sula granti*) (32, 61), waved albatrosses (*Phoebastria irrorata*) (63), Galápagos penguins (*Spheniscus mendiculus*) (64), and Galápagos petrels (*Pterodroma phaeopygia*) (65, 66). The speciation processes underlying this concentration of endemism in the eastern Pacific Ocean are largely unknown, but our findings suggest that periods of glacial maxima affect seabird distributions in this region. As evidenced by behavioral studies of the Cocos and brown boobies, assortative mating is not preempted by genomic differentiation (37). The morphological differences between Cocos and brown boobies as well as their tendency for assortative mating suggest that some barriers to gene flow have emerged between these populations, despite high rates of hybridization (37). These patterns emphasize the importance of continued investigation of booby speciation as a model for understanding how oceanic biodiversity arises in this ecologically important region.

### Genomic Differentiation Between Recently Diverged Species Pairs

We investigated the two pairs of sister species that demonstrate evidence of introgression for signatures of genomic regions involved in species divergence. We used sliding window F_ST_ scans implemented in ANGSD and identified 13 genes in outlying regions of high F_ST_ in comparisons of blue-footed and Peruvian boobies and 76 genes in comparisons of masked and Nazca boobies (Figure 3). Fisher’s exact tests with a q-value threshold of < 0.05 identified no overrepresented GO terms in either comparison. However, investigations of all GO terms associated with one or more genes, unfiltered by p-value, identified a suite of candidate genes involved in morphological and physiological processes. Several GO terms associated with animal organ morphogenesis and skeletal system development emerged in comparisons of blue-footed and Peruvian boobies (Figure S5). Genes within genomic regions that differ between sister species were involved in many processes, with body development among the most interpretable. Blue-footed and Peruvian boobies demonstrate a few clear peaks in F_ST_, two of which contain protocadherin (PCDH) genes (Figure 3B). Protocadherins are expressed in feather bud tissues and likely determine several components of feather development (67), but they are also involved in neural development (68), and the phenotypic effects of these differentiated regions in blue-footed and Peruvian boobies are unclear. A broad suite of GO terms emerged in comparisons of masked and Nazca boobies, including phospholipid biosynthesis, carbohydrate metabolism, and response to nutrient levels (Figure S5). Genomes of masked and Nazca boobies were more diffusely differentiated, with outlying F_ST_ windows spread throughout the genome. This is potentially due to our global sampling scheme of masked boobies, which led to high variation within the masked booby samples. Given our evidence of high diversity across pantropical species, investigations with deep sampling in individual localities are likely to be more fruitful in uncovering genes underlying species differences in boobies.

### Evolution of the Neo-Sex Chromosome

We surveyed the neo-sex chromosome for gene content and putative species-specific structural differences that could influence speciation processes. We first identified northern gannet chromosome 29 as orthologous to the neo-sex chromosome in boobies by testing for significant differences in heterozygosity between male and female samples. Across the genome, we only observed higher heterozygosity in females compared to males – a signal associated with misalignment from sex chromosomes with a ZZ reference genome – on the Z chromosome and on chromosome 29 (Figure 4; Wilcoxon rank-sum test; Z chromosome: female mean H = 0.43, male mean H = 0.04, FDR corrected p < 0.001; chromosome 29: female mean H = 0.29, male mean H = 0.054, FDR corrected p < 0.001).

**Figure 4.**
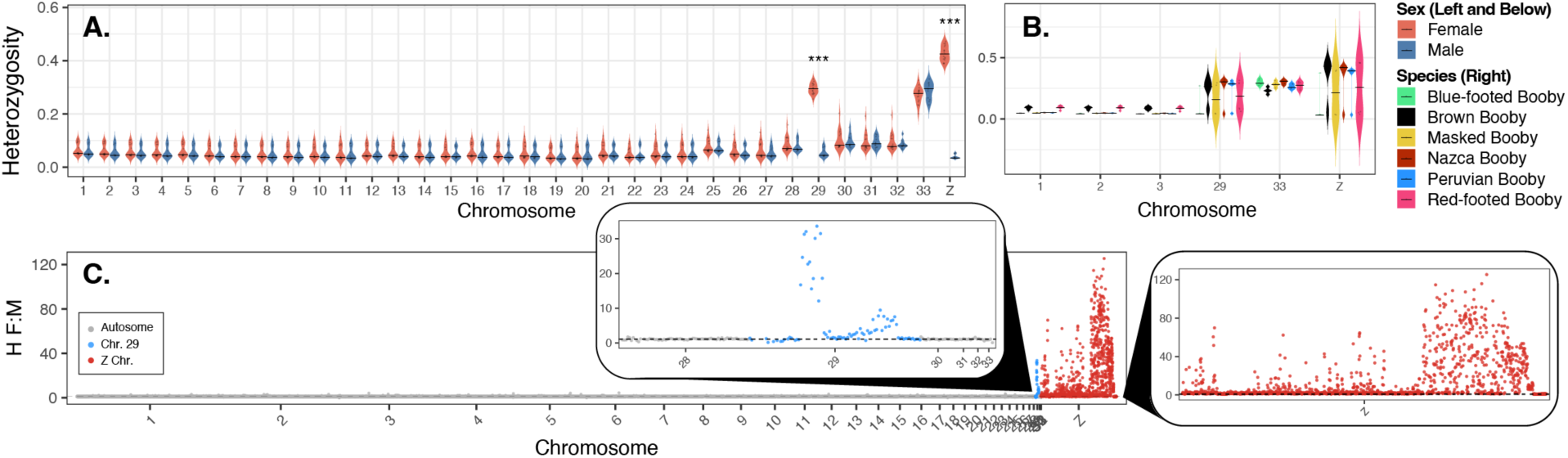
Sex inference and identification of the neo-sex chromosome. **A.** Mean heterozygosity of individuals across the genome by sex. Females exhibit higher heterozygosity than males on chromosome 29 and the Z chromosome. Both sexes demonstrate high heterozygosity on chromosome 33. **B.** Mean heterozygosity of individuals by species on chromosomes of interest and the three largest autosomes. Chromosomes with significantly different heterozygosity values between the sexes are denoted with asterisks: FDR adjusted p < 0.05 (*), p < 0.01 (**), and p < 0.001 (***). Species differences do not explain the observed patterns in sex-specific heterozygosity across chromosome 29, nor do the species differ in the high levels of heterozygosity on chromosome 33. **C.** The ratio of mean female heterozygosity to mean male heterozygosity in 50kb windows across the genome, with northern gannet chromosome 29, the proposed neo-sex chromosome in boobies, and the ancestral avian Z chromosome highlighted.

Genetic divergence between sexes within a species is only possible in genomic regions that do not recombine between the Z and W chromosomes. Recombination is only suppressed between the Z and W chromosomes in female birds, which have a Z_1_Z_2_W karyotype, but will persist between the two Z_1_ and two Z_2_ chromosomes in male birds, which have a Z_1_Z_1_Z_2_Z_2_ karyotype. We inferred patterns of recombination suppression across the neo-sex region of the W chromosome using the ratio of female-to-male heterozygosity, which revealed a complex pattern of interspersed recombining and non-recombining regions. We refer to regions with significantly greater female than male heterozygosity as putatively non-recombining between the Z_2_ and the W and we refer to those with no significant difference between female and male samples in heterozygosity as putatively recombining. We identified the centromere by characterizing chromosome-wide patterns of repetitive elements using the Vertebrata database in Repeatmasker (69) and quantifying SNP density. Region 0 has very low SNP density and the highest density of repetitive elements across chromosome 29, which suggests this is likely an acrocentric centromere. Robertsonian translocations involve the fusion of two chromosomes at the centromere, often losing one chromosomal arm from each (70). Because chromosome 29 is either telocentric or highly acrocentric in the northern gannet, this would not lead to much or any gene loss. Characterizing gene loss on the W will require a high-quality chromosome level reference genome from a female booby.

Our efforts to delineate the nonrecombining from the recombining regions of the neo-W chromosome suggest independent recombination suppression events across species. We identified two nonadjacent blocks with, and two nonadjacent blocks without, sex-associated signatures of heterozygosity, as well as one block with interspecific variation (Figure 5A-E, S6). These patterns suggest that 1. an inversion on this chromosome predated the neo-sex chromosome evolution in the ancestral booby species and 2. the recombination boundary exhibits interspecific differences (Figure 5F-H). These regions suggest an island of reduced recombination surrounded by two recombining regions, which is inconsistent with the processes by which recombination becomes restricted across sex chromosomes. More likely, either an inversion in the northern gannet lineage or in the ancestral booby lineage underlies this pattern, allowing for the order of regions in boobies to putatively be 1, 4, 3, 2, and 5 (Figure 5G-H). Under this model of a putative inversion, recombination is restricted on one side of the chromosome (Regions 1, 3, and 4) and unrestricted on the other, with interspecific variation in the boundary between the two. Region 1 demonstrates strong evidence of no recombination, further supporting the interpretation that this edge is bound to the W chromosome in boobies. Regions 2 and 5 have no sex differences in heterozygosity but regions 3 and 4 do.

**Figure 5.**
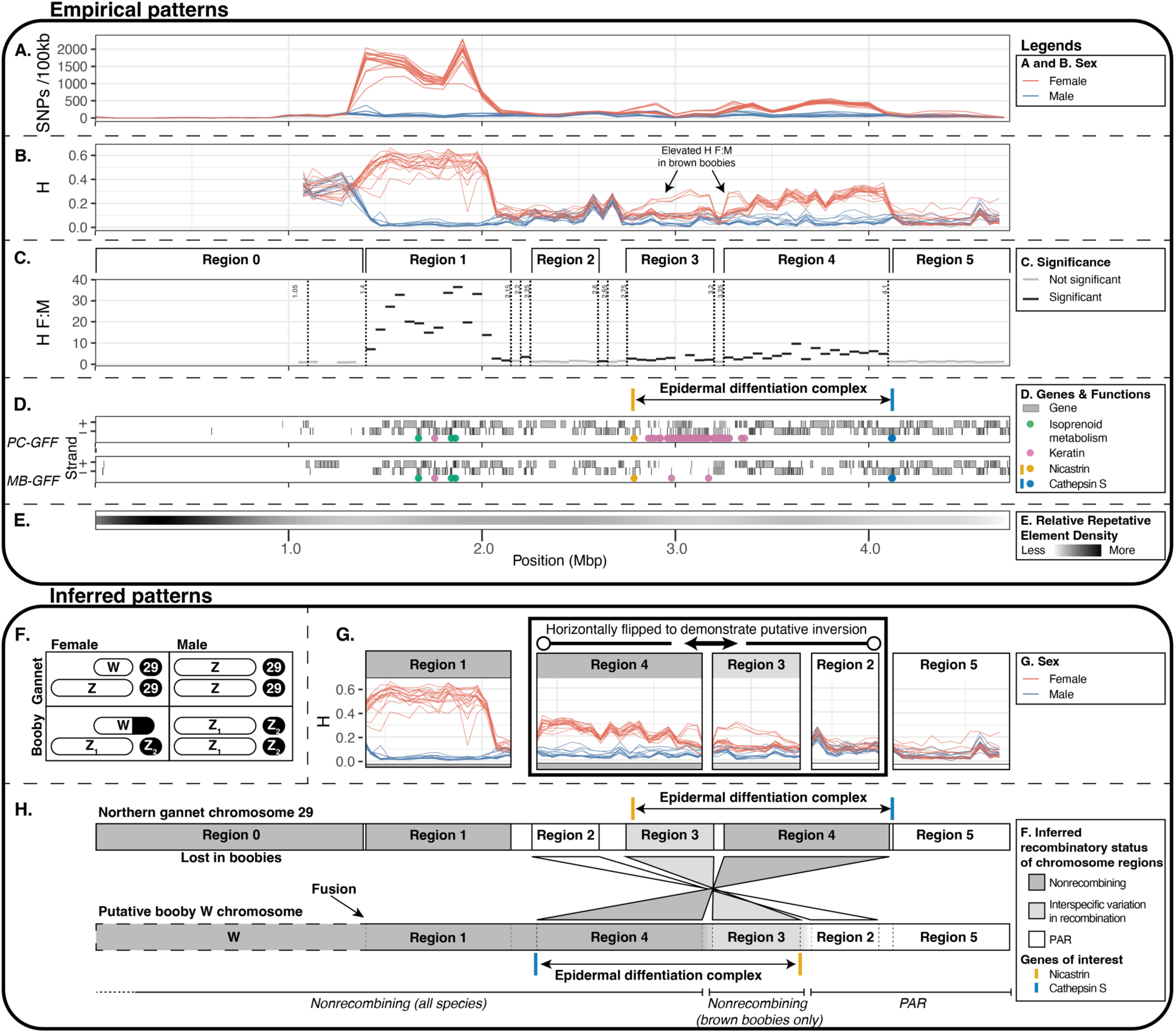
Evidence and inference of recombinational patterns in the neo-sex chromosome. Plots **A-E** depict empirical observations using the northern gannet reference genome. Plots **A-C** show statistics computed from booby genomic data that were used to infer evidence of recombination or recombination suppression across the neo-sex chromosome: Plot **A.** shows counts of SNPs in 100kb windows within each individual, plot **B.** shows individual heterozygosity in 50kb windows, and plot **C.** shows the Female:Male ratio of heterozygosity in 50kb windows, with FDR corrected significantly different windows in black (t-test, alpha of 0.001). Vertical lines depict region boundaries of consecutive runs of multiple windows of the same significance status. Plots **D** and **E** depict features of the northern gannet genome of biological relevance. Plot **D.** shows a map of all genes across the chromosome using the PC-GFF (above) and MB-GFF (below). Within each plot, the leading strand (+) is in the top row and the lagging strand (-) is in the bottom row. Genes of interest are annotated with dots of colors corresponding to the function of the gene, and the epidermal differentiation complex is annotated above. Plot **E.** shows the density of repetitive elements across the chromosome, which largely cluster within Region 0. Plot **F.** depicts karyotypes based on findings from Pozzobon et al. (28), with the addition of the northern gannet chromosome 29 label identified in our analyses. Plot **H** shows heterozygosity across the neo-sex chromosome with regions 2-4 horizontally flipped to be consistent with a putative inversion. This arrangement better explains the observed patterns of Female:Male heterozygosity across the chromosome that is consistent with the mechanisms of recombination on sex chromosomes as it results in a pattern of a high female-to-male heterozygosity ratio on the left and a low ratio on the right. Plot **G.** shows a schematic of the inferred chromosome structure in boobies, demonstrating the possible reordering of the regions on the Z2/neo-W chromosomes in boobies that could have resulted from a single inversion in the ancestral booby taxa.

Region 3 exhibits interspecific variation in the recombination boundary. This region has sex-linked heterozygosity levels in all booby species except for red-footed boobies, suggesting that a unique recombination suppression event occurred on the W chromosome after red-footed boobies split from the other taxa. However, boobies in the brown-Cocos clade exhibit much higher differences between female and male heterozygosity in this region: up to 0.2, compared to 0.09–0.10 in blue-footed, Peruvian, masked, and Nazca boobies. In blue-footed, Peruvian, masked, and Nazca boobies, recombination may persist in this region at a reduced rate, may have become restricted at a more recent time point than in brown boobies, or brown boobies may have accumulated more variation in this region of the W chromosome than other species. In the absence of de novo assembled reference genomes from each booby species, our inferences are limited, and we therefore refrain from defining unique evolutionary strata across the neo-sex region, but we identify this is a candidate for further investigation.

Interestingly, we found many genes involved in color signaling and reproductive traits that span the boundary between recombining and non-recombining neo-sex regions of the W chromosome. Given the putative nature of the inversion, we conservatively evaluated overrepresented gene ontologies within each region, rather than within inferred recombining and non-recombining regions. Isoprenoid metabolism was the only significantly overrepresented gene ontology category in a predicted non-recombining region of the neo-sex chromosome (Region 1; isoprenoid biosynthetic process, GO:0008299, q = 0.005; terpenoid biosynthetic process, GO:0016114, q = 0.014; isoprenoid metabolic process, GO:0006720, q = 0.070). This region contained 62 genes, including one involved in keratin production (KRTCAP2), three involved in isoprenoid metabolism (FDPS, PMVK, and FLAD1), and one involved in bone development (BGLAP). All three isoprenoid metabolism genes are upstream of the production of cholesterols and ubiquinones in vertebrates (71). Cholesterols are abundant in eggs and sex-specific genetic variants could be relevant to egg production. Additionally, female boobies deposit biliverdin in eggshells, an important circulating antioxidant, and other antioxidants like carotenoids and ubiquinones may be required to reduce oxidative stress during the breeding period (72, 73). Alternatively, ubiquinone supplementation increases carotenoid deposition in zebra finch bills, and these isoprenoid metabolic genes may instead function indirectly to improve carotenoid-based signals, which underlie bare-part coloration in boobies (74). These non-mutually exclusive hypotheses suggest that isoprenoid genes on the neo-sex chromosome could influence reproductive investment and color-trait signaling. In the putatively recombining regions, Region 2 contained 16 genes that were significantly associated with various GO terms including anatomical development (GO:0048856, q = 0.035; Figure S7). No GO terms were enriched in Region 5.

We also identified a suite of genes involved in the development of skin, bills, feathers, and claws in birds that spans the boundary of recombination across all boobies (Regions 2–4) called the epidermal differentiation complex (75). This complex is bounded by *nicastrin* and *cathepsin S* (*75*), which occur in Regions 3 and 4, respectively. It appears to be completely nonrecombining between the Z_2_ and W chromosomes in the brown-Cocos clade and the red-footed boobies but potentially recombines with the Z2 chromosome in the rest of the species. This region contains a cassette of keratin-related genes, cornified envelope precursors, and S100 genes that are involved in the development of the skin and skin appendages like feathers, beaks, and scales (75). Genes in regions of reduced recombination between the sex chromosomes can be involved in sexually dimorphic traits, and we found that the epidermal differentiation gene complex spans the boundary of the predicted nonrecombining and recombining regions of the W chromosome, with interspecific variation. We identified many keratin-related genes within the region of the neo-sex chromosome where recombination is uniquely reduced in brown boobies. Brown booby males and females differ in the coloration of their faces, feet, and bills (33, 36). Cocos boobies — which are genetically a lineage within brown boobies — have sexual dimorphism in feather coloration on their heads. This region is a strong candidate for the genetic mechanism underlying sex differences in coloration in brown-Cocos boobies. Part of the epidermal differentiation gene complex spans a region of predicted reduced recombination shared across all booby species. This suggests that sex-specific keratin-based traits in other booby species may also be tied to this neo-sex chromosome. Boobies are broadly sexually monomorphic in most traits, but subtle differences between the sexes do occur in some species and colonies (33). Notably, Nazca boobies have greater sexual dimorphism than masked boobies in bill morphometrics (35), another keratin-based trait. This finding implicates the neo-sex chromosome as a strong candidate region involved in the evolution of color and other epidermal traits in boobies.

Much remains to be explored in this neo-sex chromosome, including characterizing gene loss and intraspecific variation. Additionally, the lack of a W chromosome in the reference genome limited our ability to characterize sequence variation to any of the three sex chromosomes, W, Z_1_, and Z_2_, due to misalignment of the W to the Zs. An annotated booby reference genome from a female individual will greatly improve our understanding of evolution on the sex chromosomes. Species-specific variants in the epidermal differentiation complex could underlie the diversity of bare-part colors observed in boobies. This neo-sex chromosome appears to be unique to boobies, but there are bare part coloration patches in other members of the Sulidae family, including subtle eye ring coloration in gannets and foot, bill, and face coloration in Abbott’s booby (*Papasula abbotti*), which is the most anciently diverged lineage within the *Sulidae* – despite the shared name, boobies are more closely related to gannets than they are to Abbott’s boobies (50). Much more work is needed to understand how the evolution of the neo-sex chromosome has influenced booby phenotypic diversity.

Sex-linked genomic regions typically experience less introgression than autosomes (44), and structural sex chromosomal variation can contribute to reproductive isolation (76, 77). Whether and how the neo-sex region contributes to speciation in boobies is an exciting avenue for future research. Our characterization of the neo-sex chromosome also identified a putative inversion that could have facilitated speciation between boobies and gannets, although long-read genomic data from multiple booby species will be necessary to confidently define it. Boundaries of recombination on the sex-limited chromosome can and often do vary between populations within a species, and while we found no evidence of intraspecific variation, our sampling was limited with respect to this question. Future genomic investigations of boobies will certainly lead to better insight into within-species diversity.

## Conclusions

Our analyses of sister-species booby genomes support a model of speciation across alternating periods of gene flow and differentiation driven by global climate trends. We showed that historic glacial periods were associated with periods of genetic divergence between sister taxa, but that the speciation process was also characterized by periods of gene flow. We present genomic evidence that supported the species status of Nazca boobies but that indicated the Cocos booby is phylogenetically nested within brown boobies. And finally, our findings implicated a neo-sex chromosome as a putatively important genomic feature for speciation in the presence of gene flow. Our analyses of genomic booby sequences reveal complex genomic and evolutionary mechanisms underlying diversification in a widespread tropical and sub-tropical seabird clade.

## Methods

### Northern gannet genome assembly

High molecular weight DNA was isolated from a muscle tissue sample from a male northern gannet in the care of the Leibniz Institute for Zoo and Wildlife Research using a MagAttract HMW DNA Kit (Qiagen, Cat. No. 67563) and an input of 14 mg muscle. 6 µg of DNA was sheared using a Megaruptor 3 instrument (Diagenode, Denville, NJ, USA). A PacBio library was prepared with sheared DNA, a SMRTbell Prep Kit 3.0 (Pacific Biosciences, Cat. No. 102-141-700), and a Pippin HT instrument (Sage Science, Beverly, MA, USA). The library was sequenced on a PacBio Sequel IIe instrument to generate 49 Gb HiFi data. 125 Gb HiC data was generated by first cross-linking 14 mg muscle and then preparing an Arima HiC library with Arima v2 Hi-C kit (PN: A410110) and then sequenced on a Novaseq 6000 instrument with 2x150 bp read length. The genome was assembled using the VGP assembly pipeline v2.0 in Galaxy as described in Lariviere et al. 2024. Briefly, the assembly used Hifiasm-HiC (v. 0.19.3) and yahs (v. 1.2a.2), resulting in two complete haplotypes.

### Booby sampling, DNA extraction, and sequencing

We extracted DNA from blood samples from 34 birds that were sampled previously for population genetic analyses (Figure 1; Friesen et al. 2002, Morris-Pocock et al. 2011, 2016, Steeves et al. 2003, 2005a, 2005b, Taylor et al. 2010a, 2011a, 2011b) and were archived at Queens University. We sequenced our samples in 2019 but in 2024 the eastern Pacific population of brown boobies was elevated to species status, now called the Cocos booby (36, 78), from which we had only sequenced one individual. Remaining samples form a longitudinal transect of much of the range of each pantropical booby species, excluding the Indian Ocean in brown boobies (Figure 1). Sequencing was unsuccessful for one masked booby from the eastern Pacific Ocean (masked booby 3), and so this sample was excluded from all analyses.

We performed DNA extractions at the University of Colorado using a Qiagen DNeasy Blood & Tissue kit and a Thermo Scientific GeneJET Genomic DNA Purification Kit. We measured DNA concentrations on a Thermofisher Qubit 3.0, and we used a Zymo Research DNA Clean & Concentrator kit to concentrate samples with low DNA concentrations. Libraries were prepared using the Nextera XT V2 library kit and were sequenced at the Genomics and Microarray Core at the University of Colorado Anschutz Medical Campus with an Illumina NovaSEQ 6000 with a Paired End 150 cycle 2x150. Raw sequence data are publicly available for download through the Sequence Read Archive (BioProject accession: PRJNA836623).

### Data cleaning

We trimmed and analyzed raw sequence fasta files for quality using Trimmomatic v0.39 (79) and FastQC v0.11.9 (80). All analyses were performed using reads aligned to a northern gannet genome (GCA_031468815.1). This genome is a chromosome level assembly from the sister genus of boobies.

We then generated an annotation file with EGAPx (https://github.com/ncbi/egapx) using published RNA-seq data from a *Morus bassanus* individual of unknown sex (81) (Sample: SAMN13755240, SRA Accession: SRX7523318), and we refer to this throughout the text as MB-GFF. We supplemented this annotation file with LiftOn using the GFF from the closest annotated relative, the great cormorant (*Phalacrocorax carbo*, GCA_963921805.1) using the - copies flag; we refer to this as the PC-lifton-GFF. We performed genome-wide analyses of genes using only the MB-GFF. To visualize genes on the neo-sex chromosome, we used both gff files because we could independently verify relevant genes by comparing the gene order of regions of interest to orthologous genes in the chicken (*Gallus gallus*) genome. For genome-wide analyses and gene ontology (GO) overrepresentation analyses of the neo-sex chromosome, we used the more conservative MB-GFF. To infer the location of the centromere in the neo-sex chromosome, we annotated repetitive elements using the Vertebrata database in Repeatmasker (69).

We aligned trimmed reads to the reference genome using bwa mem v0.7.18, sorted bam files using SAMtools v1.19.2 (82), and then added or replaced read groups and marked duplicates using picard tools v2.23.4 (83). We then clipped overlapping reads using the clipOverlap command from bamutil v1.0.15 (84) and realigned indels using the GenomeAnalysisTK.jar function from GATK v3.7 using java version jdk1.8.0_411 (85).

We measured average depth and quality using ANGSD v0.94 (86) and excluded four individuals with less than 3X mean coverage and one with extremely high standard deviation in coverage (Table S1). We used two SNP filtering approaches, as appropriate for the analyses and pipelines. First, we ran analyses for sex assignment, principal component analyses (PCAs), and F_ST_ in ANGSD using genotype likelihoods under the SAMtools model (-GL 1). We kept only sites with a base quality of ≥ 30, mapping quality of ≥ 30, MAF ≥ 0.05, read depth ≥ 4, individual count of ≥ 15, and SNP p-value threshold of 1 × 10⁻⁶. Second, for neo-sex chromosome analysis, phylogenetic and historical demographic inference, and D-statistics, we used BCFtools v1.9 (87). We used Phred-scaled likelihoods implemented in Dsuite v0.5 to compute D-statistics (88). We used BCFtools to call variants from bam files, ignoring indels, and to filter SNPs for quality >30 and depth between 4-20x. We used plink v2.0 to keep only biallelic SNPs with <2% missing genotypes and ≥1% minor allele frequency, and to remove individuals with >20% of genotypes (89). For phylogenetic analyses, VCFs were converted to phylip files using the script vcf2phylip.py3 (90) and invariant sites were removed using the script ascbias.py (91).

### Sex inference

To infer the sex of each individual, we compared normalized depth and mean heterozygosity across chromosomes, a commonly applied and validated approach for sex inference from genomic data (92, 93). We expected to observe lower average depth on the Z in females (Z_1_Z_2_W individuals) compared to autosomal read depth. However, because the northern gannet reference genome lacks a W chromosome, we expect reads from the W to misalign to the Z_1_ and Z_2_ chromosomes, leading to higher rates of apparent heterozygosity in females. We inferred that individuals with elevated heterozygosity and reduced depth on the northern gannet Z chromosome were female (Z_1_Z_2_W) and that individuals with comparable depth and heterozygosity to the autosomes were male (Z_1_Z_1_Z_2_Z_2_). Depth scores were noisier than heterozygosity, and so we relied on heterozygosity for final interpretations (Figures S8 and S1). We sampled at least one individual inferred to be of each sex from each species, aside from the Cocos booby which had only one sampled individual, which was inferred to be female (Table S1).

### Neo-sex chromosome inference and analysis

To identify which microchromosome in the northern gannet is orthologous to the neo-sex chromosome in boobies (fused with the W chromosome but not the Z, forming a Z_2_, as identified in a previous karyotype (28)), we evaluated normalized mean depth and mean heterozygosity across all chromosomes. Normalized mean depth was noisy and provided no sex-association across any chromosome other than the Z_1_ (chromosome Z in northern gannets), so we relied exclusively on heterozygosity to infer the identity of the Z_2_. To infer regions of reduced recombination between the sex chromosomes, we computed heterozygosity and SNP density across the Z_1_ and Z_2_ chromosomes in nonoverlapping 50kb windows per individual, skipping windows with less than 100 SNPs. We tested for significant differences in heterozygosity between male and female samples per window using an FDR-corrected t-test with a q-value cutoff of 0.05. We defined a “region” as an instance of three or more continuous windows of the same significance value. Regions of significant difference were designated as putative nonrecombining regions.

To infer the biological processes affected by genes on the neo-sex chromosome, we curated gene lists within the putative recombining and nonrecombining regions and analyzed them for overrepresented Gene Ontology terms using PANTHER (FDR-corrected Fisher’s Exact Test, PANTHER GO-Slim Biological Process annotation dataset) with the genome-wide gene list as the background (94). We also manually inspected the lists for genes that could contribute to sex differences in phenotypic traits.

### Phylogenetic analysis and genome comparisons

We generated multiple species alignments of all individuals using phylip files generated from the VCF. We analyzed only autosomal chromosomes for all phylogenetic analyses, and we removed chromosome 33 given indications of poor alignment, likely resulting from the high GC content of this microchromosome (63.5% compared to a genome average of 42.5%), which is very high even for a microchromosome (95). Neither sex nor species were associated with heterozygosity on this microchromosome, so we inferred this pattern to derive from read misalignment, and we excluded this chromosome from all analyses.

We inferred phylogenetic relationships across species and individuals by running raxml-ng on each individual chromosome with a GTR+G model using a Lewis correction for ascertainment bias and generated 200 bootstraps for each analysis (96). Trees were plotted in R with phangorn (v2.12.1), ape (v5.8.1), ggtree (v3.14.0), and treeio (v1.30.0) packages (97–100). We constructed a consensus network using the consensusNet function from the ape package (97).

We analyzed whole genome clustering across samples using principal component analyses (PCA) and ADMIXTURE, implemented in PCAngsd v1.36.4 (49). PCA and ADMIXTURE analyses were performed on all individuals, between each pair of sister species, and within each species.

### Population history

We modeled historic effective population sizes of each sample using MSMC2 (101). We used SNPable to identify and filter out sites with low callability using the SNPable pipeline (seqbility v.20091110), phased VCFs of each chromosome for each individual using WhatsHap v2.8 (102), generated input using the generate_multihetsep.py function from msmc-tools, and ran MSMC2 using all autosomes of each individual as input and using the default fixed time segment pattern of 1*2+25*1+1*2+1*3. We plotted the historical pattern of effective population sizes using the estimated mutation rate of 1.91e^-9^ from chicken (103), and a range of generation times from 8–18 years with a mean of 13 years. Estimated generation lengths for boobies range from 8.5–17.3 years (104). Relative cross-coalescence rates (RCCR) were inferred by running an MSMC2 model of the combined two populations of interest and then generating a combined MSMC output file using the combineCrossCoal.py script from msmc-tools. RCCRs were plotted alongside average change in global surface temperatures which were modeled by Snyder 2016 (105).

### Testing for introgression

We employed Dsuite to compute f-branch (*f_b_*) and Patterson’s *D*-statistic using Phred-scaled genotype likelihoods (88). We evaluated the standard p-value associated with the D-statistic as well as the clustering-robust p-value designed to test for significant clustering of putatively introgressed sites (“ABBA” sites) (106). The clustering test reduces false positives derived from mutation rate variation across the tree. Two p-values are generated in Dsuite to test for clustering: the clustering-sensitive p-value and the clustering-robust. We report both in the supplement (Table S2) but restrict our interpretations to the clustering-robust value as the clustering-sensitive value is more prone to false positives (106). For each set of p-values, we used a false-discovery rate (FDR) correction for multiple testing using the Benjamini-Hochberg procedure (107).

### Differentiation

To evaluate genomic patterns of differentiation between booby taxa, we performed sliding window F_ST_ scans. We compared two species pairs (masked and Nazca; blue-footed and Peruvian) using a 50kb window and 10kb step in ANGSD to identify regions that may have undergone a selective sweep in one of the two compared populations. We considered the top 0.1% of windows to be outliers based on the negative log of the p-value. Our sample sizes is relatively small but sufficient for accurate estimation of F_ST_ (108).

### Biological inference of candidate gene functions

For each prioritized gene list (candidate genes in regions of high differentiation and genes in regions inferred to be introgressed), we assessed overrepresented biological pathways using GO terms in PANTHER using the chicken PANTHER GO-Biological Slim Annotation Dataset while correcting for multiple testing using FDR with a q threshold of <0.05. A single gene can have a major impact on species phenotype, so we additionally report any GO term associated with one or more candidate genes identified from the selection analyses. We visualized all subsets of GO terms with scatter plots of semantic dissimilarity in rrvgo (109).

## Data sharing plan

Raw sequence data are available for download from the NCBI Sequence Read Archive (SRA submission SUB11340555; will be made public prior to publication). All scripts used to analyze this dataset are available on GitHub and will be archived permanently upon acceptance (https://github.com/dannyjackson/SulidaeGenomics).

## Funding information

Genomic sequencing was funded by University of Colorado Boulder start-up funds for the Taylor Lab. An open access license was not selected.

## Acknowledgements

We would like to thank all the members of the Taylor Lab at the University of Colorado Boulder, McGraw Lab at Arizona State University, and Comparative Genomics and Reproductive Health Section in the National Human Genome Research Institute at the National Institutes of Health for their thoughtful comments on the development of this analysis and manuscript. We also thank everyone who helped to collect and process the samples for previous studies, including M. Amin Ordoñez, J. Anderson, J. Awkerman, L. L. Baglietto, P. Baião, B. Baker, R. J. Balbín, M. Betts, T. Birt, R. Both, L. Braun, J.A. Castillo-Guerrero, M. Chaib, A.C. Dávila, G. Dell’Omo, C. Depkin, K. Feeley, E. Flint, E. Flores, M. Guevara,, L. Hayes, A. Hebshi, S. Hebshi, J. Hennicke, K. Huyvaert, J. Jahncke, C. Jones, L. Liu, H. Lormee, L. Lougheed, T. Loxton, G. Luna-Jorquera,, M. Miller, M. Müller, G. Mori, A. T. Nieto, D. O’Daniel, P. O’Neill, M. Amin Ordoñez, L. Ott, P. Parker, E. Porter, N. Ratcliffe, G. Rocamora, M. Reque, S. Schaffer, M. Schiffler, Y. Sherman, A. Simeone,, R. White, J. Zamudio, and to all field assistants for their assistance in field collecting and sample processing. Samples were collected for previous work, and we thank all the agencies and organizations who helped facilitate these collections, who are impractical to list here due to space limitations but who can be found in the relevant papers (1–7). This work was supported, in part, by the Intramural Research Program of the National Human Genome Research Institute, National Institutes of Health. The contributions of the NIH authors are considered Works of the United States Government. The findings and conclusions presented in this paper are those of the authors and do not necessarily reflect the views of the NIH or the U.S. Department of Health and Human Services.

## Extended Data Figures

**Figure S1.**
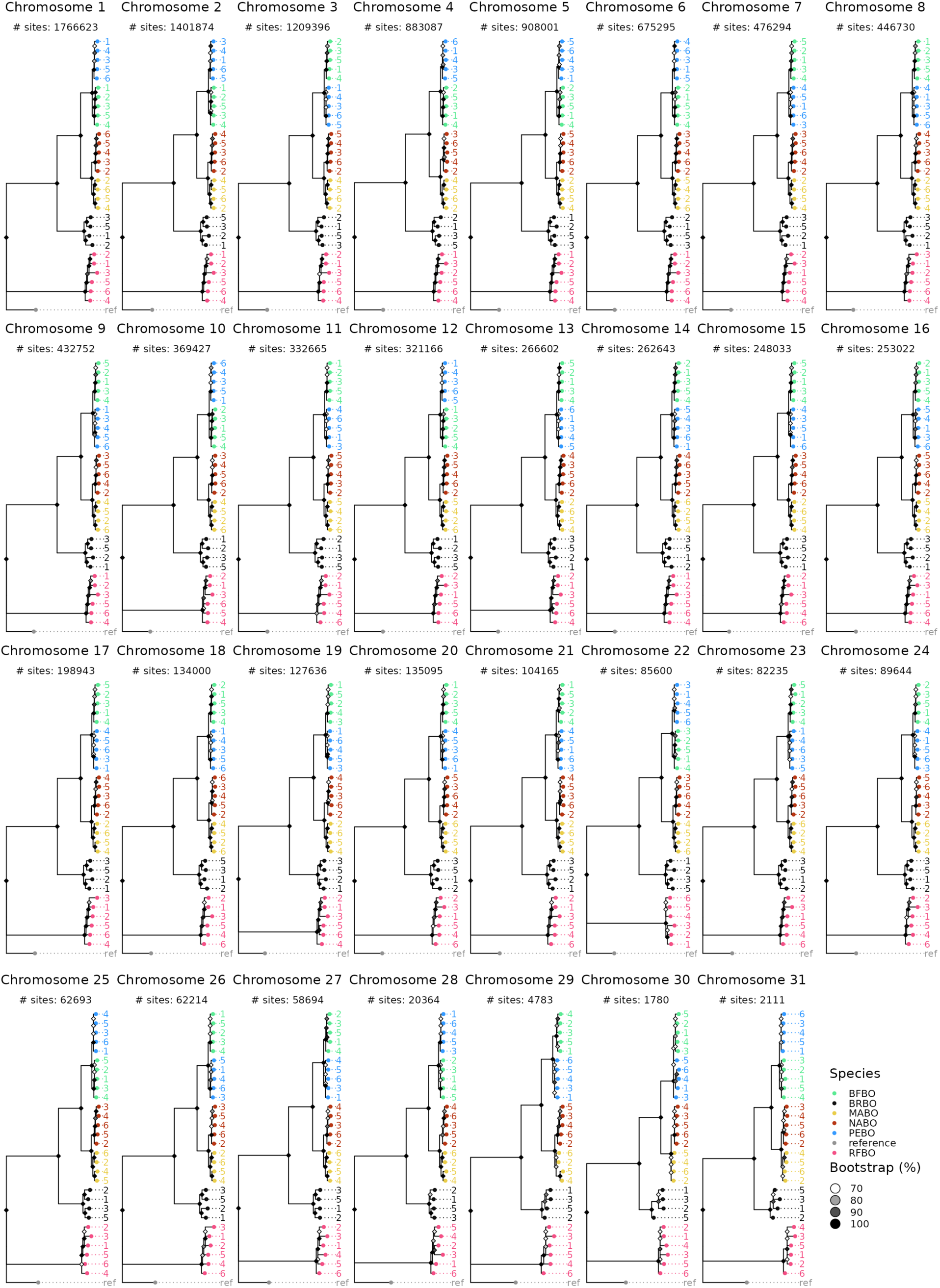
Phylogenetic trees constructed from each chromosome. Chromosome 29 was dropped from the final consensus tree after it was identified as a neo-sex chromosome. Species level monophyly was recovered across analyses of all macrochromosomes except for the relationship between brown and Cocos boobies, for which our analyses recover a deeper split in the brown-Cocos clade between ocean basins than between species.

**Figure S2.**
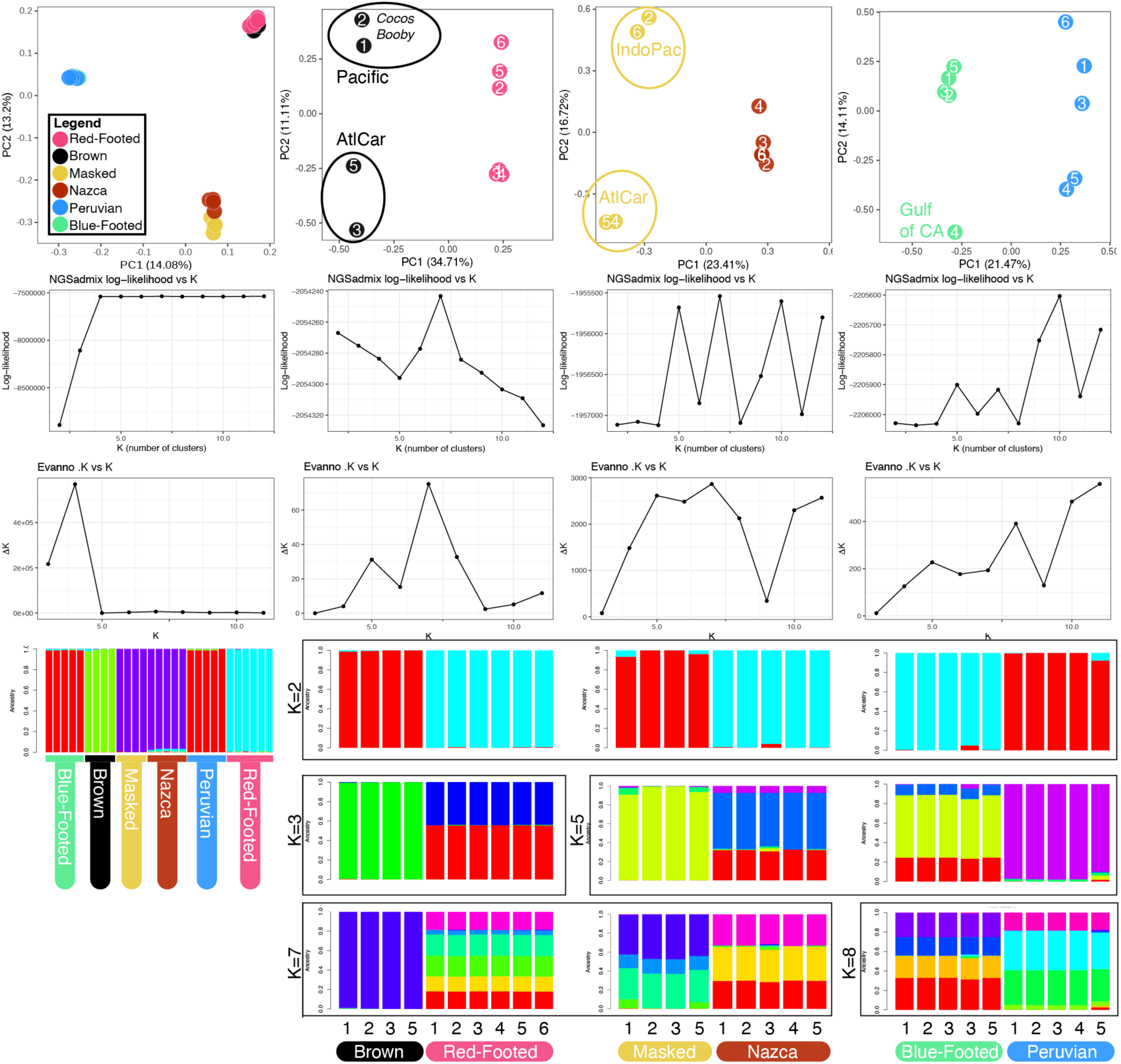
Results of principal component and ADMIXTURE analyses of all species and of species pairs. The ellipses were manually added to denote sampling locations of clusters that arise from specific ocean basins. Sample sizes are small for within-species analyses, especially for ADMIXTURE analyses, and we therefore weight insights from PCAs over those from ADMIXTURE.

**Figure S3.**
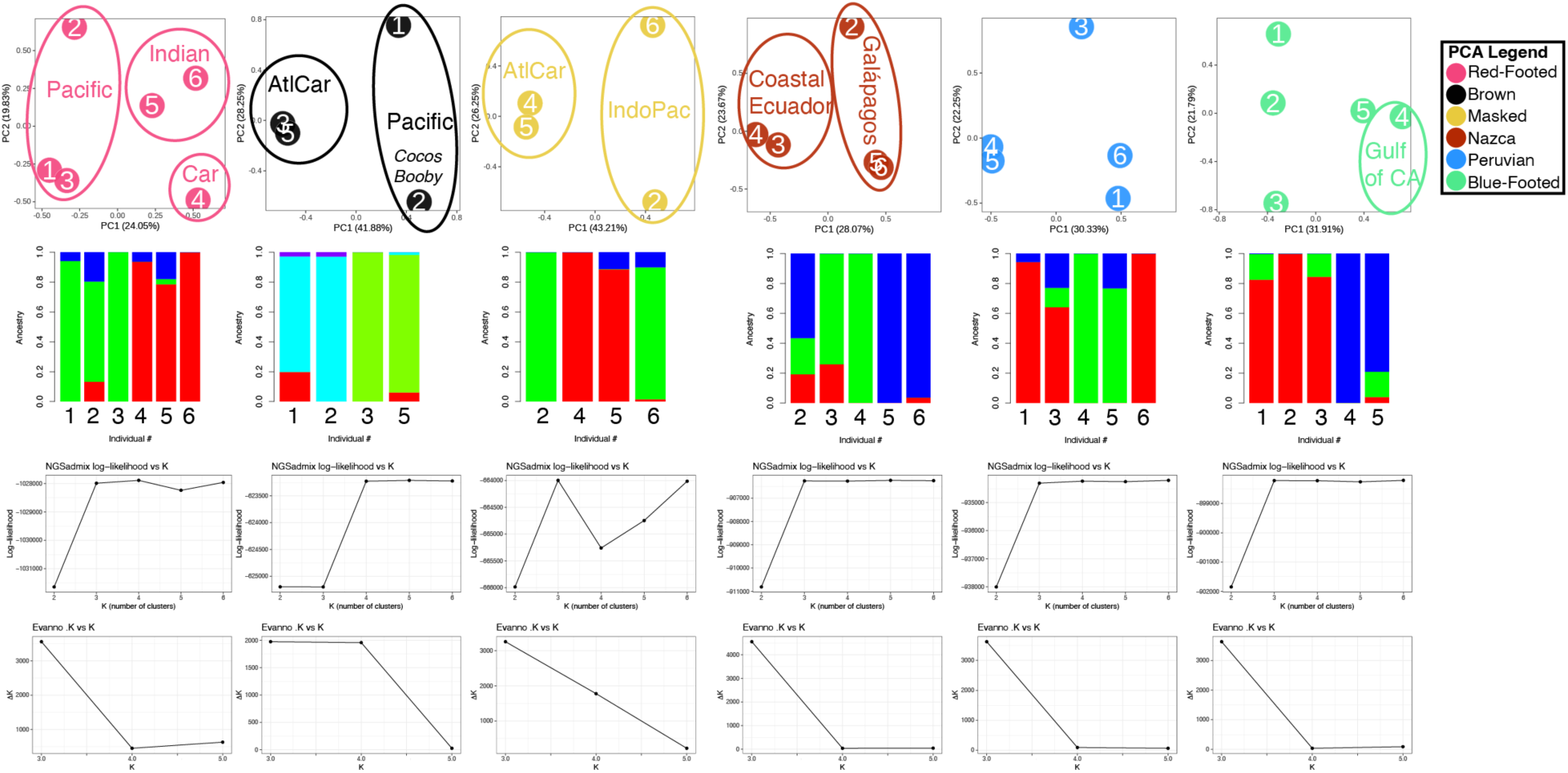
Results of principal component and ADMIXTURE analyses of populations. The ellipses were manually added to denote sampling locations of clusters that arise from specific ocean basins. Sample sizes are small for within-species analyses, especially for ADMIXTURE analyses, and we therefore weight insights from PCAs over those from ADMIXTURE.

**Figure S4.**
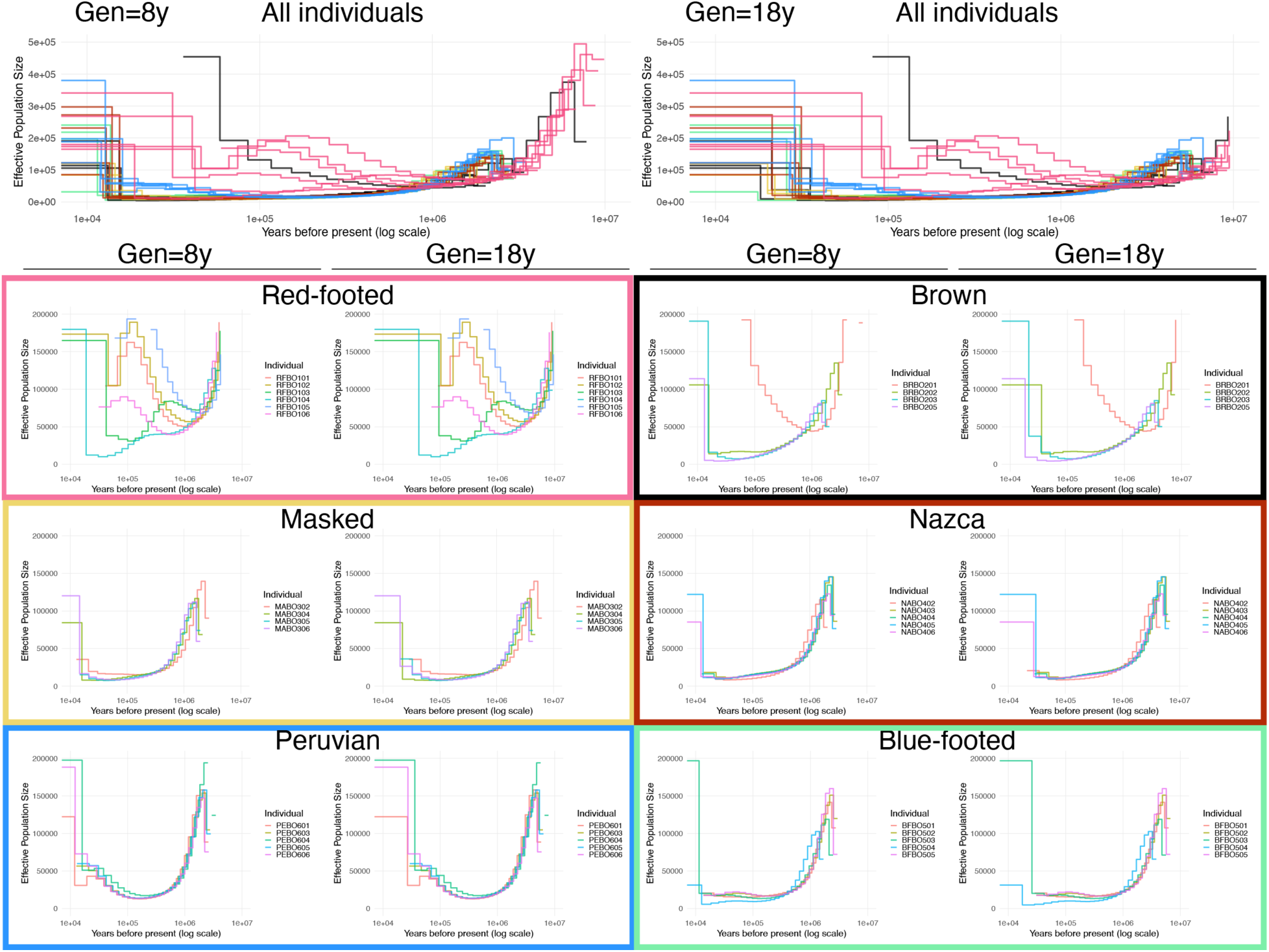
Demographic history models by individual in MSMC. Plots show the lower and upper bound generation times and are plotted colored by species (top two plots) and by individual (lower six plots). The color of the lines in the “All individuals” plots correspond to the box around each species. Historical demographic modeling of individual genomes was consistent within taxa except for red-footed and brown-Cocos boobies (Figure S6). All species showed a sharply declining trend prior to one million years ago, and masked, Nazca, Peruvian, and blue-footed boobies remained at that low effective population size over time. Genomes from different populations of red-footed boobies varied over the last million years, with the samples from the eastern Pacific and Caribbean declining in recent years and all other samples increasing. The central Pacific brown booby sample is an outlier from other brown booby samples, and this genome demonstrated population size expansion over the last million years, while other brown booby samples had patterns of genetic variation consistent with declining effective population size. The Cocos booby, central Pacific masked booby samples, and equatorial blue-footed booby samples showed a slightly higher effective population size compared to their conspecifics (or for the Cocos booby, compared to brown boobies) around 10,000–100,000 years ago. All Peruvian booby samples increased in effective population around that time as well. The Gulf of California blue-footed booby sample demonstrated a lower effective population size than the equatorial blue-footed booby population over the last ∼100,000 years.

**Figure S5:**
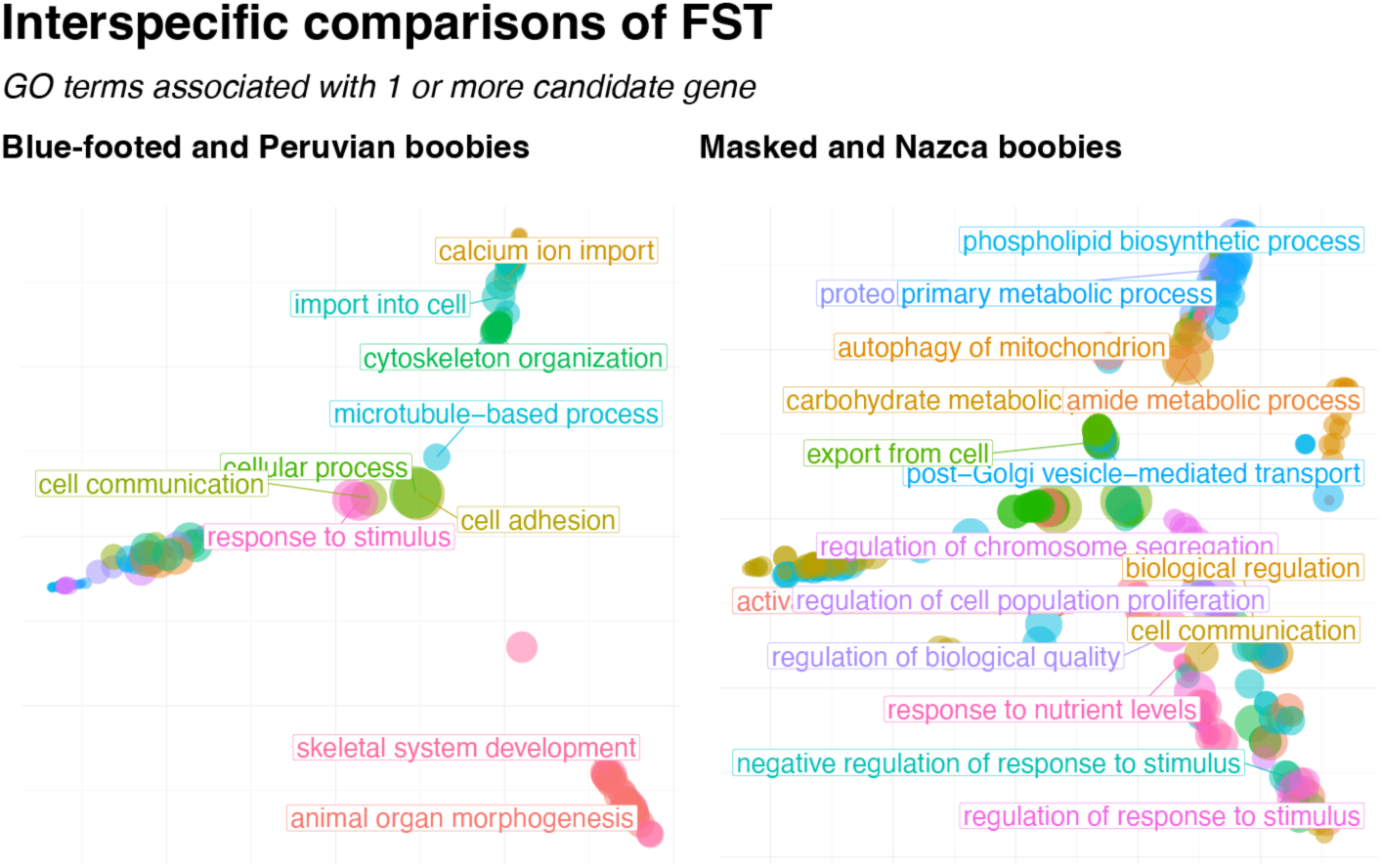
Semantic dissimilarity plots of enriched GO terms in outlier gene lists from interspecific F_ST_ comparisons.

**Figure S6.**
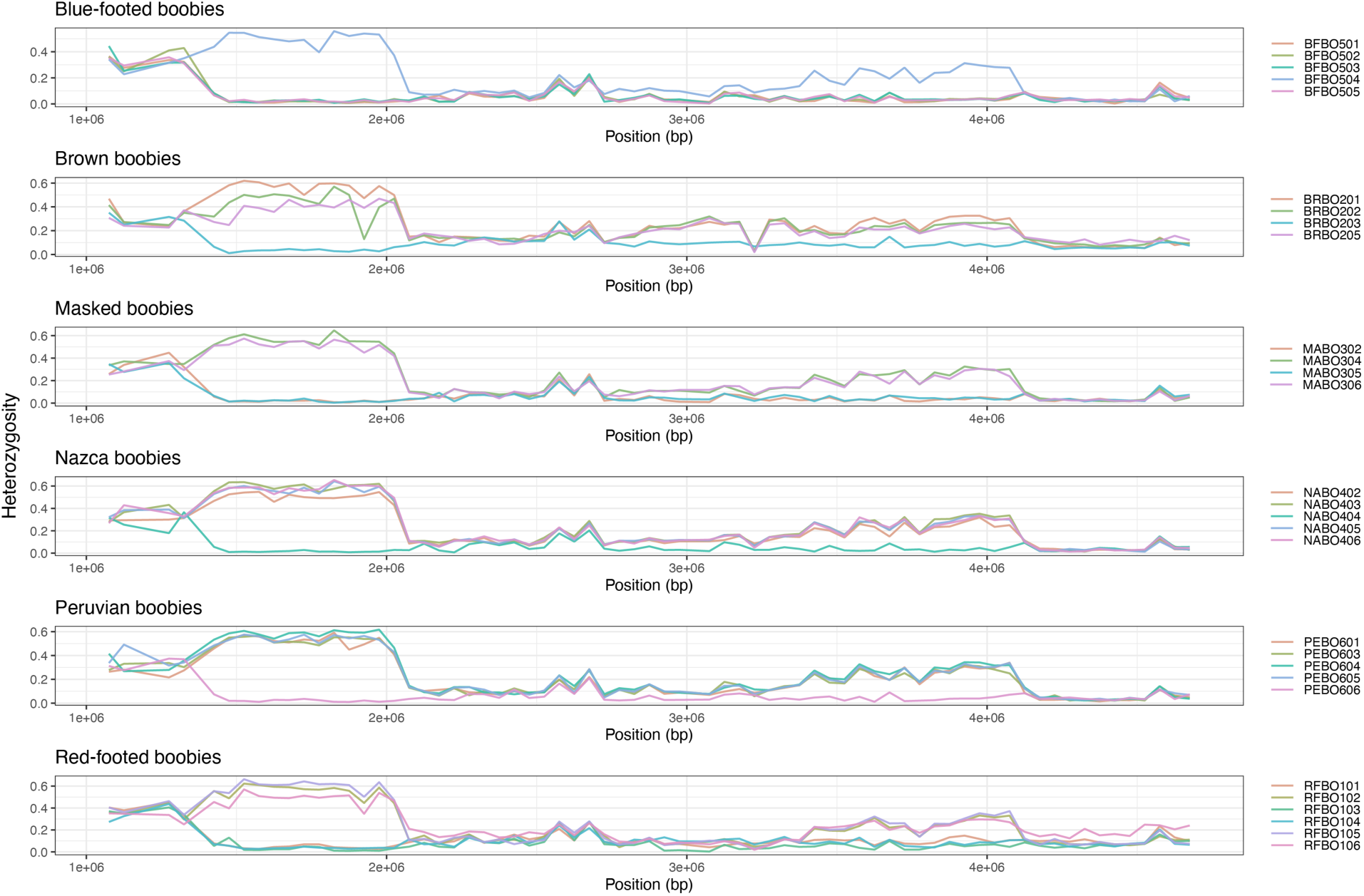
Heterozygosity across the neo-sex chromosome, faceted by species and colored by individual.

**Figure S7:**
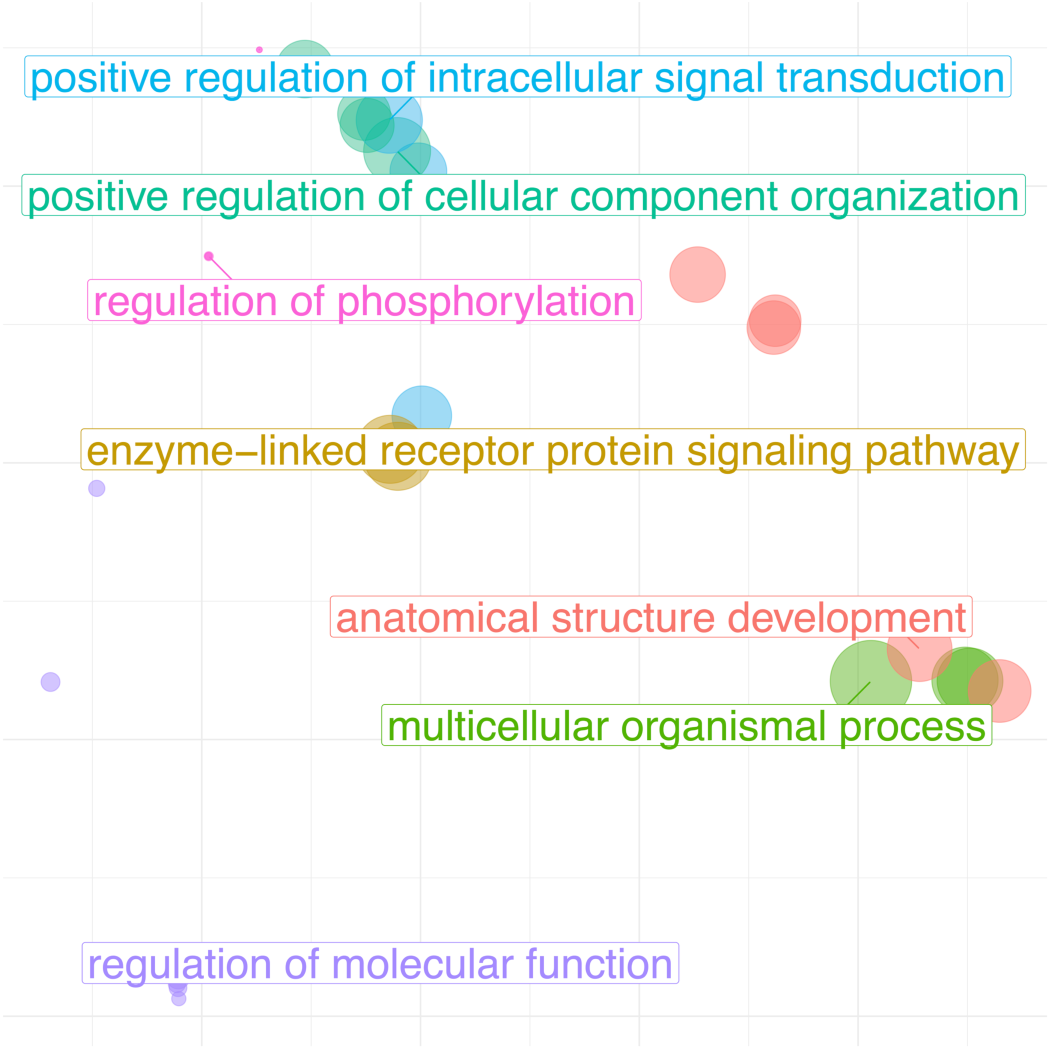
Semantic dissimilarity plot of enriched GO terms within region 2 of the neo-sex chromosome.

**Figure S8:**
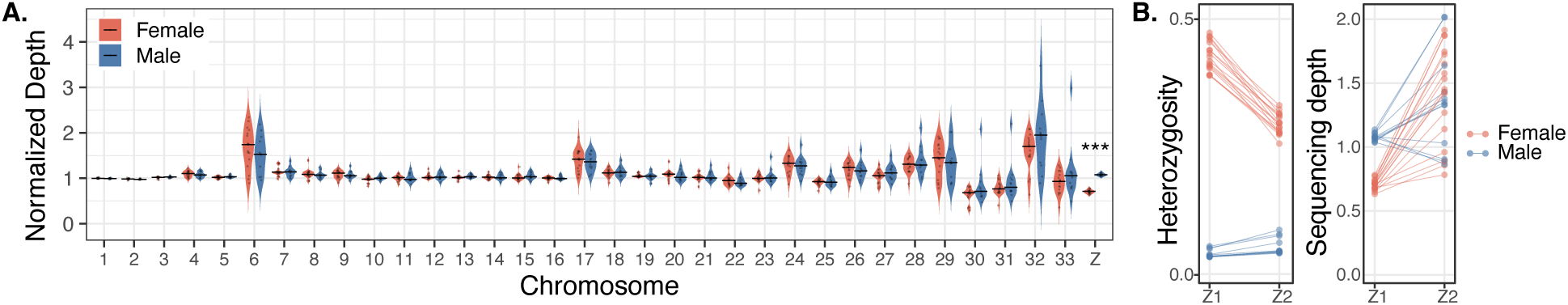
**A**. Depth of samples by chromosome and by inferred sex. The Z chromosome is the only chromosome with significant differences in depth between the sexes (Wilcoxon rank-sum test; Z chromosome: female mean H = XXX, male mean H = XXX, FDR corrected p < 0.001). **B**. Comparisons of heterozygosity and sequencing depth for inferring sex (using Z1) and inferring the identity of the neo-sex chromosome (Z2).

## Supplemental Tables

**Table S1.** Depth of coverage. Depths were calculated from the final bam files, after all filtering efforts, but not including any downstream filtering steps made on the VCFs. Italicized individuals had an average depth below our cutoff of 3x and were excluded from all further analyses. We excluded blue-footed booby 6, brown booby 4, masked booby 1, Nazca booby 1, and Peruvian booby 2 from all analyses based on depth statistics. Values on the basis by which an individual was excluded from future analyses are highlighted in red.

| **Individual** | **Avg. Depth** | **SD Depth** | **Inferred sex** |
| --- | --- | --- | --- |
| Blue footed 1 | 5.13 | 35.00 | Male |
| Blue footed 2 | 5.39 | 31.24 | Male |
| Blue footed 3 | 4.67 | 26.28 | Male |
| Blue footed 4 | 3.90 | 27.92 | Female |
| Blue footed 5 | 5.40 | 31.55 | Male |
| Blue footed 6 | 2.41 | 42.58 | NA |
| Brown 1 | 5.73 | 26.09 | Female |
| Brown 2 (Cocos) | 5.24 | 32.52 | Female |
| Brown 3 | 4.01 | 28.16 | Male |
| Brown 4 | 3.20 | 162.60 | NA |
| Brown 5 | 3.73 | 37.39 | Female |
| Masked 1 | 2.90 | 19.41 | NA |
| Masked 2 | 5.43 | 18.77 | Male |
| Masked 4 | 4.45 | 32.36 | Female |
| Masked 5 | 4.13 | 32.67 | Male |
| Masked 6 | 4.11 | 25.15 | Female |
| Nazca 1 | 2.58 | 26.27 | NA |
| Nazca 2 | 3.92 | 29.61 | Female |
| Nazca 3 | 5.52 | 25.45 | Female |
| Nazca 4 | 5.53 | 22.58 | Male |
| Nazca 5 | 5.44 | 29.53 | Female |
| Nazca 6 | 5.12 | 29.81 | Female |
| Peruvian 1 | 4.15 | 18.06 | Female |
| Peruvian 2 | 2.59 | 25.18 | NA |
| Peruvian 3 | 4.41 | 36.39 | Female |
| Peruvian 4 | 5.19 | 36.95 | Female |
| Peruvian 5 | 4.38 | 35.38 | Female |
| Peruvian 6 | 4.46 | 31.57 | Male |
| Red footed 1 | 6.44 | 20.53 | Male |
| Red footed 2 | 7.43 | 22.59 | Female |
| Red footed 3 | 13.85 | 40.19 | Male |
| Red footed 4 | 4.94 | 14.66 | Male |
| Red footed 5 | 13.23 | 37.29 | Female |
| Red footed 6 | 5.27 | 20.12 | Female |
| Average | 5.12 | 32.70 | 12 males |
| Avg. After exclusions | 5.34 | 28.82 | 17 females |

**Table S2:** Significant tests of introgression using D-statistics with clustering-sensitive and FDR corrected p-values.

| P1 | P2 | P3 | D statistic | CS FDR<br>q value | CR FDR<br>q value |
| --- | --- | --- | --- | --- | --- |
| Gulf of CA blue footed | Equatorial blue footed | Atlantic/Caribbean brown | 0.071 | 0.000 | 1.000 |
| Gulf of CA blue footed | Equatorial blue footed | Pacific brown | 0.046 | 0.000 | 1.000 |
| Gulf of CA blue footed | Equatorial blue footed | Atlantic/Caribbean masked | 0.150 | 0.000 | 1.000 |
| Gulf of CA blue footed | Equatorial blue footed | Indopacific masked | 0.153 | 0.000 | 1.000 |
| Gulf of CA blue footed | Equatorial blue footed | Nazca | 0.140 | 0.000 | 1.000 |
| Gulf of CA blue footed | Equatorial blue footed | Peruvian | 0.223 | 0.000 | 1.000 |
| Pacific brown | Atlantic/Caribbean<br>brown | Gulf of CA blue footed | 0.250 | 0.000 | 1.000 |
| Gulf of CA blue footed | Atlantic/Caribbean<br>masked | Atlantic/Caribbean brown | 0.039 | 0.008 | 1.000 |
| Gulf of CA blue footed | Indopacific masked | Atlantic/Caribbean brown | 0.044 | 0.007 | 1.000 |
| Gulf of CA blue footed | Nazca | Atlantic/Caribbean brown | 0.034 | 0.000 | 1.000 |
| Gulf of CA blue footed | Peruvian | Atlantic/Caribbean brown | 0.060 | 0.001 | 1.000 |
| Gulf of CA blue footed | Atlantic/Caribbean<br>masked | Pacific brown | 0.030 | 0.004 | 1.000 |
| Gulf of CA blue footed | Indopacific masked | Pacific brown | 0.035 | 0.003 | 1.000 |
| Gulf of CA blue footed | Nazca | Pacific brown | 0.031 | 0.001 | 1.000 |
| Gulf of CA blue footed | Peruvian | Pacific brown | 0.054 | 0.003 | 1.000 |
| Atlantic/Caribbean masked | Indopacific masked | Gulf of CA blue footed | 0.013 | 0.001 | 1.000 |
| Atlantic/Caribbean masked | Nazca | Gulf of CA blue footed | 0.004 | 0.001 | 1.000 |
| Gulf of CA blue footed | Peruvian | Atlantic/Caribbean masked | 0.129 | 0.000 | 1.000 |
| Nazca | Indopacific masked | Gulf of CA blue footed | 0.007 | 0.002 | 1.000 |
| Gulf of CA blue footed | Peruvian | Indopacific masked | 0.130 | 0.000 | 1.000 |
| Gulf of CA blue footed | Peruvian | Nazca | 0.128 | 0.000 | 1.000 |
| Pacific brown | Atlantic/Caribbean<br>brown | Equatorial blue footed | 0.269 | 0.000 | 1.000 |
| Atlantic/Caribbean masked | Equatorial blue footed | Atlantic/Caribbean brown | 0.012 | 0.000 | 1.000 |
| Indopacific masked | Equatorial blue footed | Atlantic/Caribbean brown | 0.006 | 0.000 | 1.000 |
| Nazca | Equatorial blue footed | Atlantic/Caribbean brown | 0.017 | 0.000 | 1.000 |
| Peruvian | Equatorial blue footed | Atlantic/Caribbean brown | 0.008 | 0.267 | 1.000 |
| Atlantic/Caribbean masked | Equatorial blue footed | Pacific brown | 0.002 | 0.001 | 1.000 |
| Equatorial blue footed | Indopacific masked | Pacific brown | 0.003 | 0.001 | 1.000 |
| Nazca | Equatorial blue footed | Pacific brown | 0.000 | 0.003 | 1.000 |
| Equatorial blue footed | Peruvian | Pacific brown | 0.014 | 0.131 | 1.000 |
| Atlantic/Caribbean masked | Indopacific masked | Equatorial blue footed | 0.016 | 0.000 | 1.000 |
| Nazca | Atlantic/Caribbean masked | Equatorial blue footed | 0.005 | 0.004 | 1.000 |
| Peruvian | Equatorial blue footed | Atlantic/Caribbean masked | 0.011 | 0.000 | 1.000 |
| Nazca | Indopacific masked | Equatorial blue footed | 0.020 | 0.018 | 1.000 |
| Peruvian | Equatorial blue footed | Indopacific masked | 0.014 | 0.001 | 1.000 |
| Peruvian | Equatorial blue footed | Nazca | 0.000 | 0.001 | 1.000 |
| Pacific brown | Atlantic/Caribbean brown | Atlantic/Caribbean masked | 0.259 | 0.000 | 1.000 |
| Pacific brown | Atlantic/Caribbean brown | Indopacific masked | 0.260 | 0.000 | 1.000 |
| Pacific brown | Atlantic/Caribbean brown | Nazca | 0.251 | 0.000 | 1.000 |
| Pacific brown | Atlantic/Caribbean brown | Peruvian | 0.254 | 0.000 | 1.000 |
| Atlantic/Caribbean masked | Indopacific masked | Atlantic/Caribbean brown | 0.008 | 0.013 | 1.000 |
| Nazca | Atlantic/Caribbean masked | Atlantic/Caribbean brown | 0.007 | 0.424 | 1.000 |
| Atlantic/Caribbean masked | Peruvian | Atlantic/Caribbean brown | 0.006 | 0.000 | 1.000 |
| Nazca | Indopacific masked | Atlantic/Caribbean brown | 0.015 | 0.133 | 1.000 |
| Indopacific masked | Peruvian | Atlantic/Caribbean brown | 0.001 | 0.000 | 1.000 |
| Nazca | Peruvian | Atlantic/Caribbean brown | 0.011 | 0.000 | 1.000 |
| Atlantic/Caribbean masked | Indopacific masked | Pacific brown | 0.007 | 0.005 | 1.000 |
| Atlantic/Caribbean masked | Nazca | Pacific brown | 0.002 | 0.051 | 1.000 |
| Atlantic/Caribbean masked | Peruvian | Pacific brown | 0.012 | 0.005 | 1.000 |
| Nazca | Indopacific masked | Pacific brown | 0.005 | 0.070 | 1.000 |
| Indopacific masked | Peruvian | Pacific brown | 0.007 | 0.000 | 1.000 |
| Nazca | Peruvian | Pacific brown | 0.010 | 0.009 | 1.000 |
| Atlantic/Caribbean masked | Indopacific masked | Nazca | 0.136 | 0.000 | 0.013 |
| Atlantic/Caribbean masked | Indopacific masked | Peruvian | 0.014 | 0.000 | 1.000 |
| Atlantic/Caribbean masked | Nazca | Peruvian | 0.005 | 0.000 | 1.000 |
| Nazca | Indopacific masked | Peruvian | 0.007 | 0.008 | 1.000 |

## Notes

### Competing Interest Statement

The authors have declared no competing interest.

